# Coupling Bacterial Cell Size Regulation with Clonal Proliferation Dynamics Reveals Cell Division Based on Surface Area

**DOI:** 10.1101/2023.12.26.573217

**Authors:** César Nieto, Sarah Täuber, Luisa Blöbaum, Zahra Vahdat, Alexander Grünberger, Abhyudai Singh

## Abstract

Single cells actively coordinate growth and division to regulate their size, yet how this size homeostasis at the single-cell level propagates over multiple generations to impact clonal expansion remains fundamentally unexplored. Classical *timer* models for cell proliferation (where the duration of the cell cycle is an independent variable) predict that the stochastic variation in colony size will increase over time. In stark contrast, implementing size control according to *adder* strategy (where on average a fixed size added from cell birth to division) leads to colony size variations that eventually decay to zero. While these results assume a fixed size of the colony-initiating progenitor cell, further analysis reveals that the magnitude of the intercolony variation in population number is sensitive to heterogeneity in the initial cell size. We validate these predictions by tracking the growth of isogenic microcolonies of *Corynebacterium glutamicum* in microfluidic chambers. Approximating their cell shape to a capsule, we observe that the degree of random variability in cell size is different depending on whether the cell size is quantified as per length, surface area, or volume, but size control remains an adder regardless of these size metrics. A comparison of the observed variability in the colony population with predictions suggests that proliferation matches better with a cell division based on the cell surface. In summary, our integrated mathematical-experimental approach bridges the paradigms of single-cell size regulation and clonal expansion at the population levels. This innovative approach provides elucidation of the mechanisms of size homeostasis from the stochastic dynamics of colony size for rod-shaped microbes.

## Introduction

Cell proliferation and cell size regulation are related processes. During division, cell size is halved and simultaneously the population increases once two daughter cells are generated. Although interconnected, cell size and cell population dynamics have been explored almost independently [1–3]. Stochastic properties of cell populations have been studied mainly using approaches independent of cell size [4–6]. From the perspective of cell size, recent research has revealed that most cells that grow exponentially control their division and therefore their proliferation employing the *adder* division strategy, where cells add, on average, a fixed size from cell birth to division [7, 8]. Although experiments on cell size statistics have been well documented [9–15], only a few details on the effects of cell size regulation on population expansion have been studied [16–21]. These efforts mainly focus on analyzing average quantities, such as the mean growth rate or the cycle duration, with weak consideration of its stochastic properties [22–24].

There is a substantial random variability (noise) between colonies in their population size [1, 20, 25]. The degree of population variability has notable consequences from an ecological perspective [26, 27]. For example, noisy cell proliferation can allow the fixation of a mutant strain in competition with others, even without showing a fitness advantage [28–30]. Furthermore, analyzing the growth dynamics of the microbial population can reveal details of pathogen infection in the host [31, 32] and how tumors proliferate [33]. This type of analysis can also be used in other areas, such as designing experiments to determine the rate of antibiotic survival in fluctuation tests [34, 35].

Proliferation has been explored mainly using an approach independent of cell size, revealing that population variability is proportional to cell cycle noise [19, 36, 37]. However, the use of new cell tracking techniques suggests that cell proliferation must consider additional cell division variables such as cell size and its growth rate [23, 38, 39]. One of the most popular microfluidic devices used for cell tracking is the *mother machine*, where a rod-shaped cell (the *mother* cell) is trapped in a closed channel and all its descendants are discarded once they emerge [40, 41]. Tracking the growth and division statistics of these trapped cells helped to discover the above-mentioned *adder* strategy [41, 42].

The origin of the *adder* is still an active research area, as cell division involves multiple mechanisms [13, 43]. Among them, we can highlight septum ring formation [11, 44], DNA replication [45], and cell wall synthesis [46, 47] that have to reach high synchronization in a complex way [48–50]. Instead of considering all details, recent studies consider phenomenological models of cell size regulation to obtain simple conclusions by exploring the link between cell division and cell size [10, 51–53]. These models can predict the dynamics of the cell size distribution with high precision [12, 53, 54]. Statistical moments of this distribution are known to present damped oscillations over time reaching a steady value [12, 55–58]. However, it is not clear whether these oscillations can also affect the dynamics of colony growth.

This article aims to understand the relationship between cell division, colony proliferation, and the geometric properties of cell size. We focus on the regulation of cell size in *Corynebacterium glutamicum* cells, which possess a rod-shaped morphology and are limited by cell wall synthesis, making the cell surface a crucial variable [59, 60]. Since our microfluidic device allows cells to grow with fewer shape restrictions compared to the *mother machine*, we anticipate a greater variability in cell width. We expect to observe differences when using different cell size proxies, such as length, volume, and surface area [61–63]. Additionally, since descendant cells can divide for approximately six generations before being overcrowded, our microfluidic device allows us to study the transient dynamics of colony populations at low cell numbers. We measure and compare these proliferation dynamics for simulated trends using different division strategies. Simulations predict different trends depending on the use of cell length, surface area, and volume on the simulation. We compare these simulated results with the variability of cell population in synchronized colonies concluding which cell size proxy is dominant on determining the division.

## Results

### Cell proliferation strategies: *timer* versus *adder*

In this section, we compare the dynamics of cell population variability for the *adder* and *timer* division strategies. The *timer*, a widely used model of population dynamics, assumes that cells divide on average after a fixed time [26,36,37,64,65]. In contrast, the *adder* considers that the duration of the cell cycle is related to the cell size [10,51]. First, we will describe how to study the statistics of the colony population. Then, we present the results for the models using mathematically simple approaches and show how they differ in their predictions of population variability. Finally, we explain how to modify these models to better compare them with the experimental data.

#### Quantification of the population variability

To study the dynamics of proliferation, we start each colony with a single progenitor cell. This progenitor is a newly born cell of size *s*_*b*_ (see Fig. 1A). We define the beginning of colony growth as the instant of progenitor birth (*t* = 0). Let *τ*_*d*_ be the random variable that defines the duration of the cell cycle, that is, the time between two consecutive divisions. After its cell cycle has completed, this progenitor will split into two cells that will also grow and proliferate. If we use the bracket notation ⟨.⟩ to represent the mean value, *τ*_*d*_ has a mean of ⟨*τ*_*d*_⟩, known as the doubling time [66]. Since ⟨*τ*_*d*_⟩ is a stochastic variable (is noisy), the colony population or the number of individuals in the colony, denoted by *N* will be also stochastic. To quantify how the division strategy affects the noise in *N*, we calculate the dynamics of the statistical moments of *N* estimated over different independent colonies. The first moment ⟨*N*⟩ corresponds to the mean colony population. The variability of the population is quantified using the squared coefficient of variation 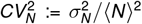, where 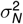 is the variance of *N*. A summary of the symbols used throughout this manuscript is provided in Table 1.

**Table 1:** Symbols used in the text.

| Variable | Interpretation |
| --- | --- |
| $N$ | Cell population in a colony |
| $CV_N^2 := \frac{\sigma_N^2}{\langle N \rangle^2}$ | Variability of the population across colonies. |
| $\tau_d$ | Cell-cycle duration. |
| $CV_{\tau_d}^2 := \frac{\sigma_{\tau_d}^2}{\langle \tau_d \rangle^2}$ | Noise in the cycle duration. |
| $s$ | Cell size (with a general metric). |
| $\mu$ | Exponential cell growth rate. |
| $s_b$ | Newborn cell size (also progenitor). |
| $s_d := s_b e^{\mu \tau_d}$ | Size at division. |
| $\Delta_d := s_d - s_b$ | Added size before division. |
| $CV_{\Delta_d}^2 := \frac{\sigma_{\Delta_d}^2}{\langle \Delta_d \rangle^2}$ | Noise in the added size before division. |
| $CV_{s_b}^2 := \frac{\sigma_{s_b}^2}{\langle s_b \rangle^2}$ | Noise in the progenitor cell size. |
| $V$ | Cell volume. |
| $A$ | Cell surface area. |
| $L$ | Cell length. |
| $w$ | Cell effective width. |
| $A_p$ | Projected cell area. |

**Figure 1:**
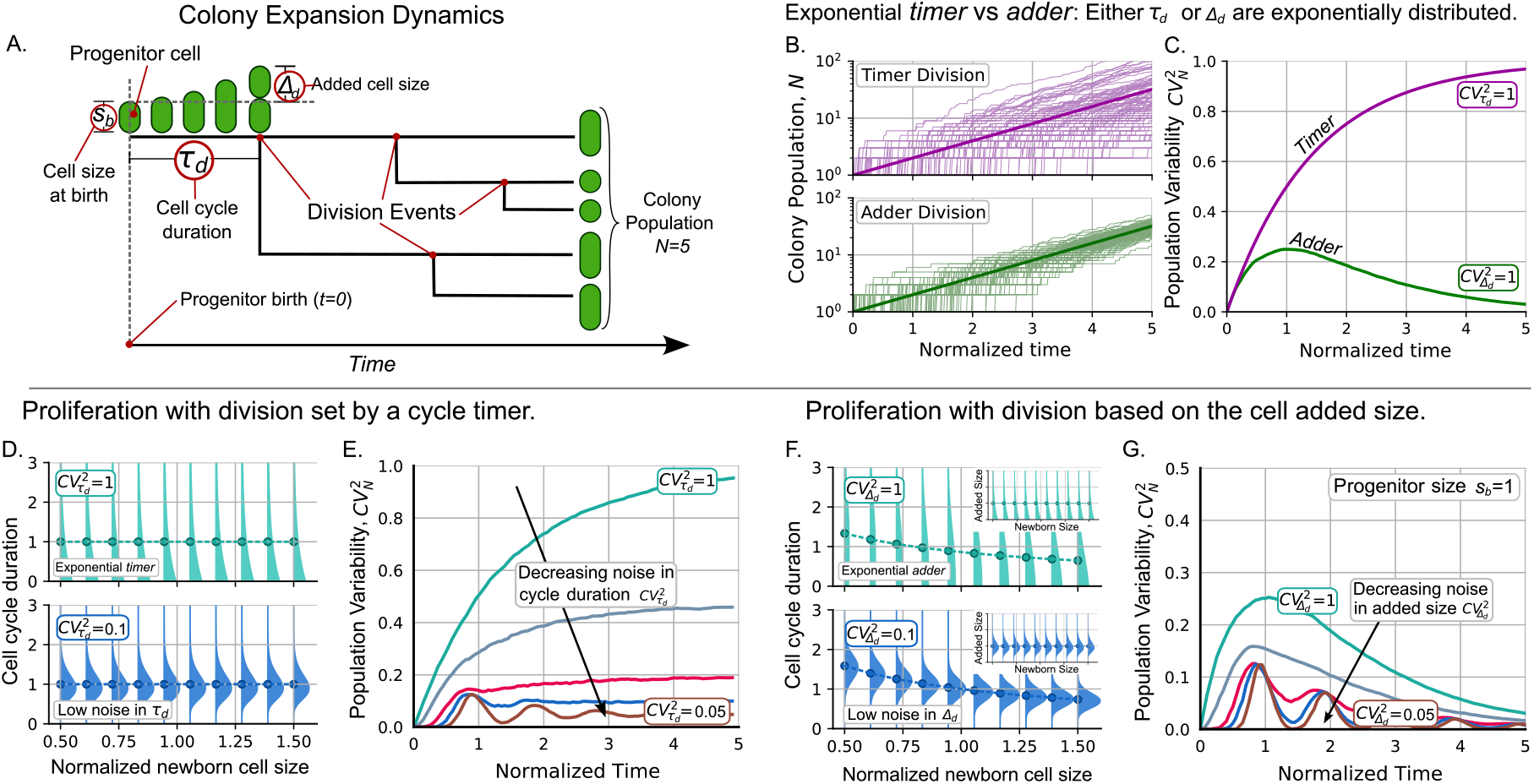
The *adder* division strategy is more effective in controlling the stochastic variation in colony population compared to the *timer* division strategy. **A**. Main variables in the colony expansion dynamics. **B**. Examples of the colony population dynamics for *timer* (top pink) and *adder* division (bottom green). **C**. Variability of the colony population over time for the two division strategies in their simplest approach. **D**. In the *timer* model, the cycle duration *τ*_*d*_ is independent and its variability is given by 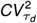. **E**. Dynamics of the variability of the colony population for 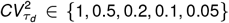. **F**. In the *adder* model, the distribution of *τ*_*d*_ depends on the size at the beginning of the cell cycle *s*_*b*_ through the added size Δ_*d*_ in (4). The mean *τ*_*d*_ given *s*_*b*_ (dots) decreases with *s*_*b*_. *Inset:* the mean added size ⟨Δ_*d*_⟩ is independent of *s*_*b*_. **G**. Dynamics of 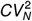 for 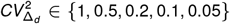. While for the *timer* 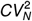 increases to a constant value, for the *adder*, 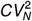 reaches its maximum around the first division and then decreases to zero. **Parameters**: For *timer, τ*_*d*_ is distributed by gamma with the specified ⟨*τ*_*d*_⟩ and 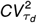. For *adder, s*_*b*_ = ⟨Δ_*d*_⟩ = 1 and Δ_*d*_ is distributed by gamma with the specified 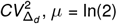.

#### Models with exponentially distributed cell cycle variables: The simplest approximation

Many theoretical models simplify cell cycle variables by assuming that they follow an exponential distribution. The exponential *timer* assumes that the duration of the entire cell cycle is distributed exponentially. For the exponential *adder* model, it is the added size during the cell cycle that follows an exponential distribution [15, 67]. Previously, we developed numerical methods to estimate population statistics over time for cells with cell cycle variables following an exponential distribution [1].

For the exponential *timer*, division occurs at a constant rate 1*/* ⟨*τ*_*d*_⟩. Thus, the division times follow an exponential distribution, which is the reason for its name. This exponential distribution has the property that the noise in the cell cycle time satisfies 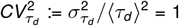 independently of ⟨*τ*_*d*_⟩. Seminal articles on cell cycle regulation [36] found that the distribution of colony population number when they proliferate following the exponential *timer* is geometric with mean ⟨*N*(*t*)⟩= exp(*µt*) with *µ* = 1*/* ⟨*τ*_*d*_⟩.

Notice that, by this formula, the population will double every ln(2) ⟨*τ*_*d*_⟩ and not every ⟨*τ*_*d*_⟩. This discrepancy between the mean cell-cycle duration and the population doubling time is due to the high noise in *τ*_*d*_, as it follows an exponential distribution [23, 36]. As the cycle duration is less noisy 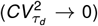, the population doubling time aligns closer with the mean cell-cycle duration [22, 23, 68]. For the upcoming results, given 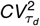, we will modify *µ* so that the population doubles every unit of time.

Given the properties of the geometric distribution for the cell population, 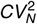 for the exponential *timer* increases over time as follows:

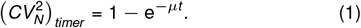

Some examples of population trajectories with cell division following the exponential *timer* are shown in Fig. 1B top. Furthermore, Fig. 1C shows how noise asymptotically satisfies 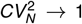 as time increases. In the general *timer*, the cell-cycle duration is drawn from a distribution less noisy than the exponential [37] and theoretical approaches have obtained useful analytical expressions [36]. The variability dynamics for this general case will be analyzed in next section.

On the other hand, for the exponential *adder* division strategy, the cell size *s* grows exponentially at a rate *µ* following the differential equation:

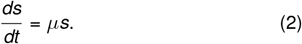

At the beginning of the cycle, we assume that the cell has random size *s*_*b*_, also known as the cell size at birth. At the end of the cell cycle, the cell has a size *s*_*d*_ called the size at division. The added size during the cell cycle is defined as the difference between the cell size at division and the size at birth Δ_*d*_ := *s*_*d*_ − *s*_*b*_. After integrating (2), we can relate these size variables to the cell-cycle duration *τ*_*d*_, as:

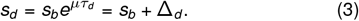

The exponential *adder* assumes that the division propensity is not constant (as the *timer*) but proportional to the cell size [10, 69]. In previous articles [51] we show how, using this rate, Δ_*d*_ is predicted to be a random variable with exponential distribution 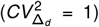 as shown in Fig. 1F. This means that *τ*_*d*_ is not an independent variable as in the *timer* mechanism but it depends on *s*_*b*_ through Δ_*d*_ following the formula obtained from (3):

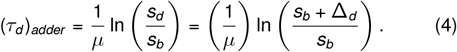

As an important property of the *adder*, by coupling cell size division and proliferation, the cell size growth rate is exactly the colony expansion rate. The doubling time in the population is exactly the mean cell-cycle duration and they are related to the growth rate by ⟨*τ*_*d*_⟩ = ln(2)*/µ* regardless of the noise in the regulation timing of division.

With these details on how, depending on the division strategy, *τ*_*d*_ or Δ_*d*_ define the division time, we will use an agent-based algorithm to obtain the population statistics. In this kind of simulation, each individual in the population will have its own attributes such as cell size, added size, and time since last cell division. During division, a new agent (a new daughter cell) is generated with its own properties. For the *timer*, the cell-cycle duration is a random variable while for the *adder*, the cell-cycle duration depends on the added size and follows (4). Details on the simulation for the *timer* and *adder* models are presented in the SI Appendix, Sections S1 and S2, respectively.

In the top panel of Fig. 1F, we illustrate how, given *s*_*b*_ and the exponential distribution of Δ_*d*_, the distribution of *τ*_*d*_ depends on *s*_*b*_. This dependency influences the dynamics of 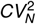 As shown in Fig. 1C, for the *adder* strategy, 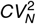 reaches its peak around the first division and then gradually decreases over time until it approaches zero. This suggests that different colonies will have asymptotically similar cell numbers, as depicted in the bottom panel of Fig. 1B.

#### Population variability is more effectively buffered by the *adder* strategy than by the *timer* strategy

In previous section, we consider models with exponentially-distributed cell-cycle variables. While the exponential *timer* has noise in cycle time 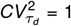, the exponential *adder* has noise in added size 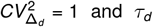 has statistics that depend on the size at birth. Experimentally, these variables are less noisy [41]. While several molecular mechanisms contribute to this precise control, we lack direct experimental data on them. Therefore, instead of building a detailed mechanistic model, we quantify the degree of noise and sample *τ*_*d*_ and Δ_*d*_ from phenomenological distributions without needing to model the particular underlying molecular detail.

In the *timer* division strategy, *τ*_*d*_ will be drawn from a gamma distribution with controlled cell cycle time 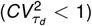. Fig. 1E shows that, similar to case 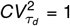, by controlling the noise in the cell cycle time, 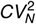 increases over time, eventually reaching a steady positive value. For high levels of control of cell-cycle duration 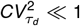 (for example, 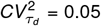) in Fig. 1E, 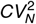 displays damped oscillations over time before eventually converging to a non-zero value. Previous research found that the asymptotical value of the fluctuations of cell population reaches a steady value proportional to the variability in *τ*_*d*_ [19, 36, 37].

For the *adder* strategy, Fig. 1G shows the trends of 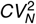 with controlled 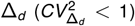. For this controlled added size, 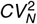 also converges asymptotically to zero similarly to the rate-based model. When the added size has noise within the typical biological range, such as 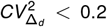 [41], 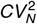 exhibits oscillations over time before converging to zero. This kind of oscillations in exponentially growing cells has been previously formally studied [70]. Our previous research found that population variability approaches to the level of variability in the colony progenitor after a long time [20].

#### Noise in progenitor cell size increases variability in colony proliferation

To plot Fig. 1G, the size of colony progenitor cell at *t* = 0 is simplified to have the deterministic value of *s*_*b*_ = ⟨Δ_*d*_⟩ with probability one. In experiments, *s*_*b*_ is a random variable with noise measured by the squared coefficient of variation 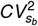. Fig. 2A shows how by increasing 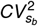 the asymptotic level of 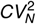 also increases. From a mathematical perspective, this means that the steady properties of the system depend on its initial conditions.

**Figure 2:**
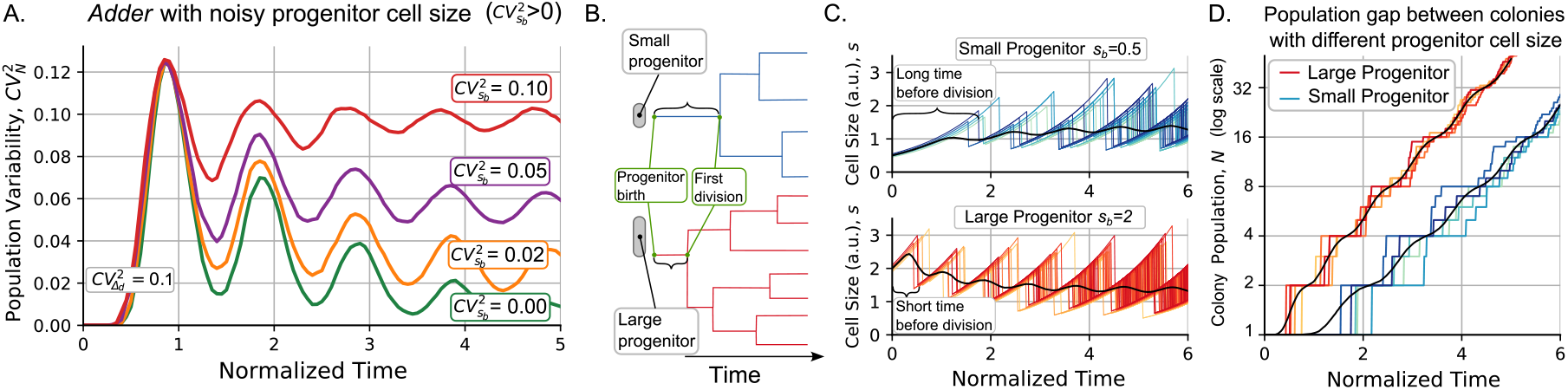
At increasing the noise in the progenitor size, the population variability does not decreases to zero but oscillates around a positive value. **A**. Dynamics of variability in the colony population 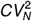 as increasing the noise in the progenitor cell size with 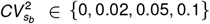. The larger the 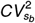, the higher the 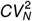 asymptotic limit, and the lower the amplitude of the oscillations. **Parameters**: 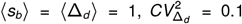. **B**. Progenitor, the first cell of the colony born at *t* = 0, may have a random size. As the division rate is proportional to cell size, larger cells will divide earlier than smaller ones. **C**. Comparison of the dynamics of colonies derived from a small progenitor (*s*_*b*_ = 0.5⟨Δ_*d*_⟩, blue shades) and a population from a large progenitor (*s*_*b*_ = 2⟨Δ_*d*_⟩, red shades). Cell size over time and **D**. Colony population over time for different colonies. Different shades represent different simulated replicas. The black lines represent the mean population, which shows a permanent gap proportional to the difference of the progenitor cell size. Observe that the y-axis has logarithmic scale.

To understand how noise in progenitor cell size influences clonal expansion dynamics, it is important to examine the relationship between a progenitor cell’s size and the timing of its first division. As illustrated in Fig. 2B, smaller progenitors typically exhibit a longer delay before their initial division compared to larger progenitors. Following this first division, the population expands asymptotically at an exponential rate (Fig. 2D). Consequently, at any given time, colonies originating from smaller cells generally have a smaller population than those originating from larger cells. This phenomenon, combined with the strong level of control that cells exert on subsequent population size once implement the *adder* to divide (Fig. 1D), implies that progenitor cell size variability is a key contributor to population variability within colonies.

An intriguing aspect of population dynamics is the oscillatory nature of their moments. Specifically, the coefficient of variation squared for the colony population 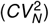 oscillates (Fig. 2A), and the mean population exhibits a staggered growth pattern, rather than a smooth exponential growth (Fig. 2D). Our results suggest that these oscillations are linked to those found in cell size moments, consistent with previous research showing that the *adder* strategy induces such cyclic behavior, as shown in Fig. 2C [12, 53, 70]. We attributed these oscillations to the coordination between cell division and growth. Here, we further show that this coordination also affects population-number statistics. In the following section, we describe experiments designed to test the periodicity of these statistics.

### Experimental setup to study the *C. glutamicum* proliferation trends

To test the validity of the predictions of our model, we study the proliferation dynamics of the nonpathogenic gram-positive soil bacterium *C. glutamicum*. Originally isolated and used due to its natural ability to excrete L-glutamate [71], *C. glutamicum* is today used for the large-scale industrial production of various amino acids, particularly L-glutamate and L-lysine [72]. At the same time, *C. glutamicum* is a well-established model species for studies related to the cell wall in *Corynebacteriales*, including prominent human pathogens such as *Mycobacterium tuberculosis* and *Corynebacterium diphtheria*, because it shares the complex organization of the cell envelope with its pathogenic relatives [73].

We used the microfluidic single-cell cultivation (MSCC) device [74, 75] to grow *C. glutamicum* cells. In this experimental setup, depicted in Fig. 3A, cells can grow and proliferate for approximately 6 generations while we maintain a controlled temperature and nutrient supply [76]. During this time, we captured snapshots of cell proliferation through phase-contrast imaging. To segment cell contours and track lineages, we use DeLTa, an image processing software [77] followed by manual verification. Additional details on the experiments and the analysis workflow can be found in Methods section.

**Figure 3:**
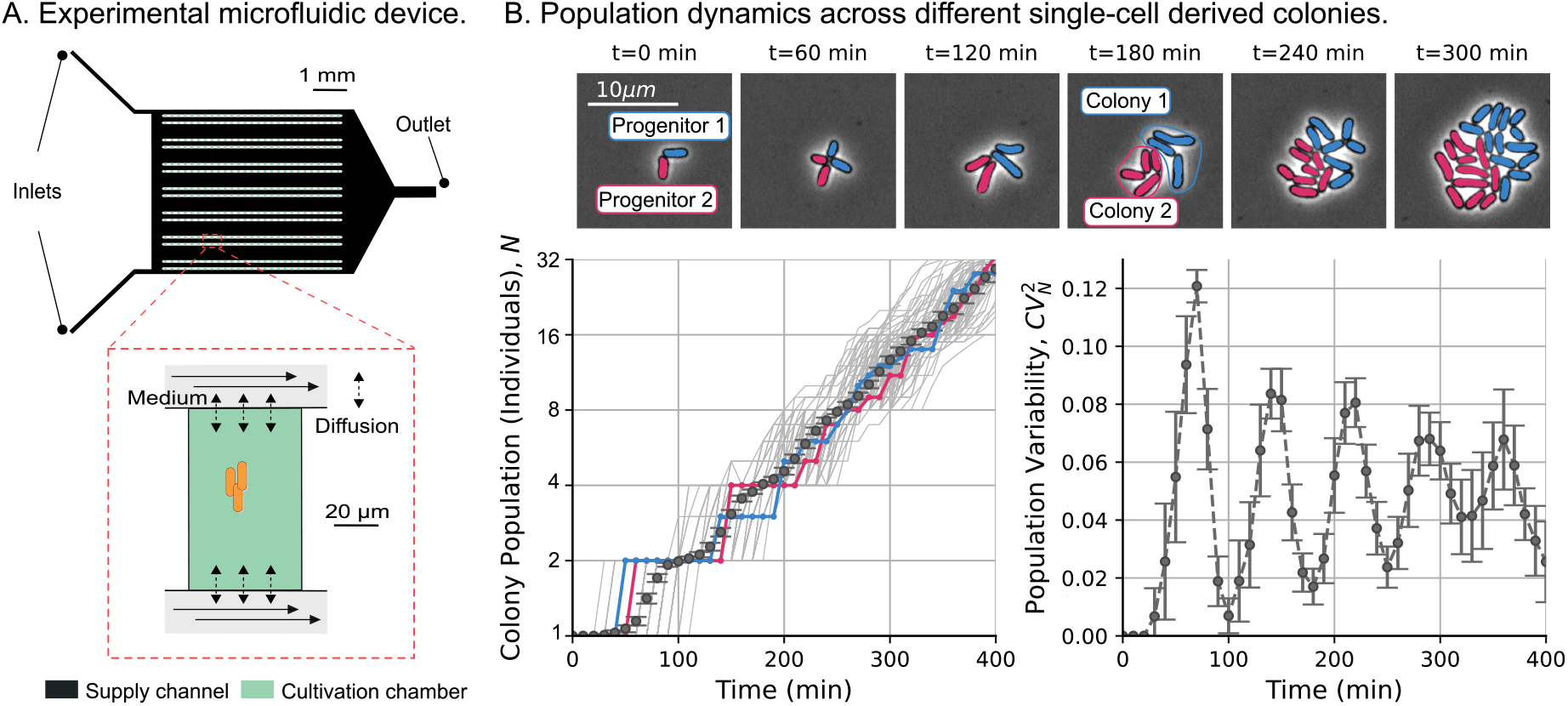
Experimental setup to investigate fluctuations in *C. glutamicum* colony population. **A**. The microfluidic single-cell cultivation (MSCC) device supplies a constant nutrient flow in 14 arrays of cultivation chambers using two inlets. **B**. (Top:) we show snapshots of the cell population for two colonies over time (*t*) each derived from a single progenitor. Different colors (blue and pink) represent different colonies. (Bottom-left:) Number of cells over time for 154 colonies (gray), with colored trajectories representing the colonies shown on top. Error-bars indicate the mean number of individuals over the studied colonies. (Bottom-right:) Noise in the colony population as the squared coefficient of variability over time (the error bars represent the 95% confidence interval for each statistical moment using bootstrapping methods).

Next, we track the growth of cell colonies. For us, a colony consists of all descendants of a given progenitor cell, and colony growth begins at the progenitor birth (Fig. 3A). This theory requires synchronization of all colonies from the birth of their respective progenitor. This synchronization is generally difficult to achieve experimentally and must be done *a posteriori* when analyzing the data [12]. In Fig. 3B we notice that the number of cells appears to grow similarly to the simulated data in Fig. 2D. Furthermore, 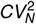 shows oscillations as predicted in Fig. 2A. Next, we will conduct a more comprehensive investigation of these oscillations and their relationship to cell size regulation.

### Geometric analysis *C. glutamicum* cell size

The main conclusion from the previous sections is that the added size noise 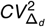 and the progenitor size noise 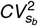 are the primary parameters to define the dynamics of the population variability in a growing colony following the *adder* division strategy. It should be noted that the particular definition of 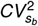 does not assume any specific geometric property for cell size. In this section, we examine how 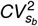 differs if cell size is measured in terms of length, surface area, or volume.

#### Sphero-cylindrical approximation to bacterial shape

To simplify the cell geometry, we approximate the cells as spherocylinders or capsules (Fig. 4A), which consists of a cylinder capped by two hemispheres [78]. We estimate the cell dimensions from the images by segmenting the cell contours and measuring the projected area *A*_*p*_, which is proportional to the pixel count within the contour (Fig. 4B). We define the cell length *L* as the longest side of the minimum-bounding rectangle of the contour. The projected area *A*_*p*_ and the cell length *L* can be related trough an *effective* cell width *w* by the formula of the projected area of a capsule:

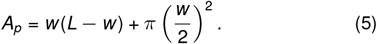

**Figure 4:**
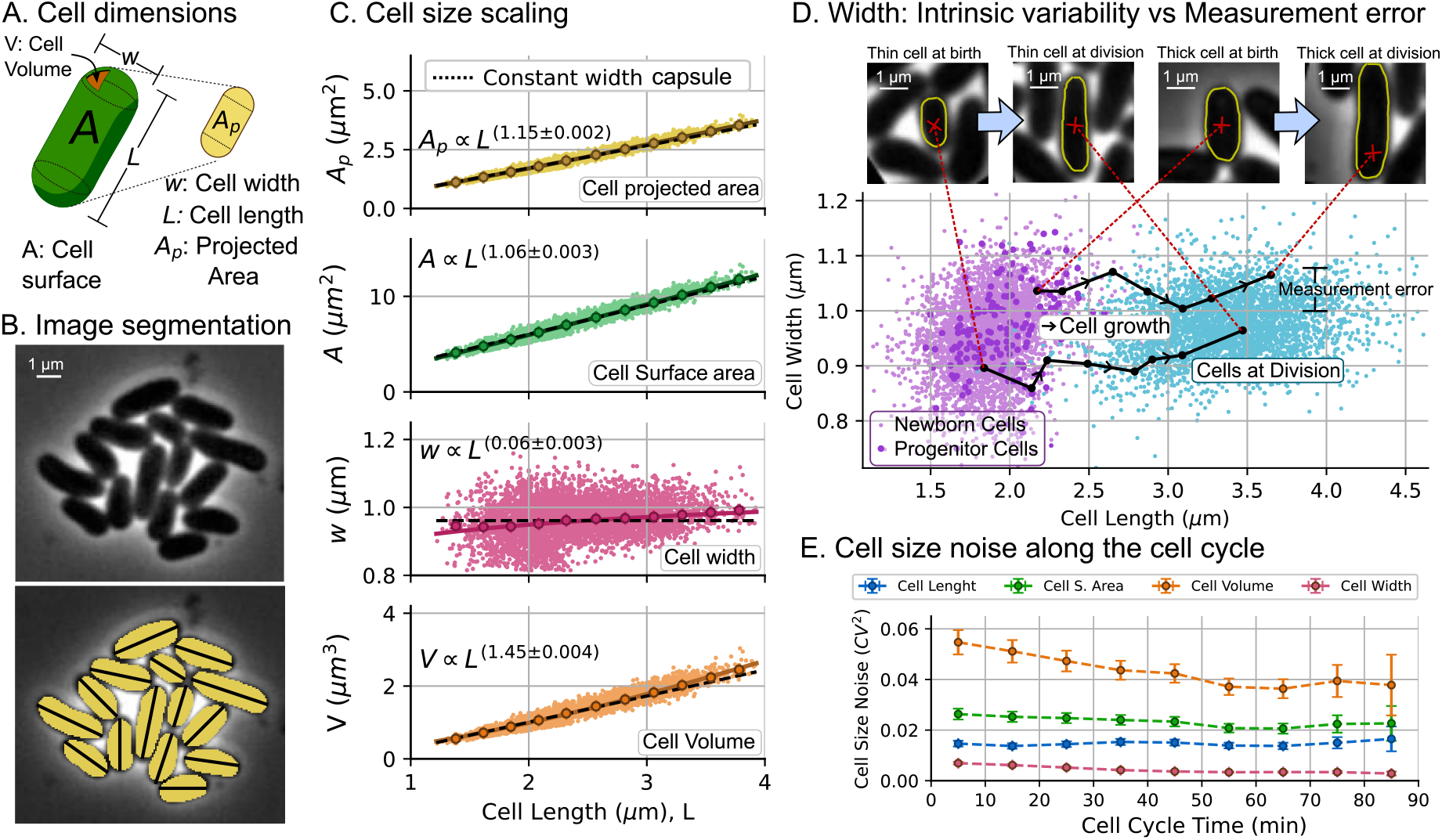
Different proxies of cell size (length, surface area, volume) yield different level of intercellular variability in cell size. **A**. The cell shape is approximated to a capsule, with projected area *A*_*p*_ and length *L* measured from the images. The width *w*, volume *V*, and surface area *A* are estimated from *A*_*p*_ and *L*. **B**. A microscopic image example with cell segmentation (yellow pixels) and measured length (black lines). **C**. Experimental data (40891 datapoints) of projected area *A*_*p*_, cell surface area *A*, cell width *w*, and cell volume *V* versus cell length *L* are compared to the approximation of the capsule shape with constant width and observed length (black dashed line). Power-law fitting (solid colored line) with the exponent is shown inset. **D**. Cell width versus cell length for newborn cells (violet) and cells at division (teal). On top, we present two cells (one thin and another thick) at the beginning and end of one cycle. The black lines show the cell dynamics along the cycle. **E**. Random variability of different cell dimensions, measured by the squared coefficient of variability as a function of the cycle time. Error bars represent the 95% confidence intervals for the statistics.

Thus, from *A*_*p*_ and *L, w* is estimated by solving (5). To illustrate the precision of this approximation for *w*, we selected two cells (Fig. 4D) with distinct and noticeable widths. The black lines in the plot width versus length in Fig. 4D represent the width inference taken for the same cells at various points in their cell cycles until division. The fluctuations of these paths give us an idea of the error associated with the shape approximation (5) (approx 3%). Cell-to-cell variability (approx 15%) is greater than this inference error.

Once estimated the cell width *w*, the cell surface area *A* and volume *V* can be estimated as follows:

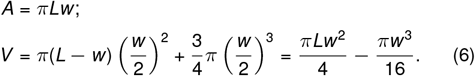

Fig. 4C illustrates how cell dimensions (*A*_*p*_, *A, w*, and *V*) scale with the length of the cell *L*. One notable observation is that area and volume do not scale perfectly linearly with cell length. Previous studies [41,79] have assumed that cell size, area, and length scale linearly, supported by the low variability of cell width in confining microfluids such as the *mother machine*. However, SI Appendix, Section S3 shows that given the assumption of a capsule shape, non-linear scaling exponents emerge naturally even if the width is constant. In fact, closer observation of Fig. 4C shows how the approximation of constant width fits the best-adjusted power laws (black dashed line). A more detailed approach about how a non-constant width can change the scaling exponent is presented in the SI Appendix, Section S3.

#### Different proxies of cell size convey different statistics

The purpose of studying the geometric properties of cell is to understand how variability in width and cell length affects the cell size noise. Consider that cells have a length with mean ⟨*L*⟩ and noise 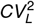; a width with mean ⟨*w*⟩ and noise 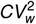. A dimension *F* (surface area, volume, etc) calculated from *w* and *L* will have a noise approximated by:

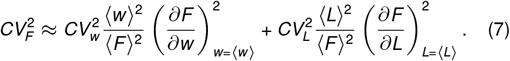

As an example of (7) for the surface area *F* = *A* := *πLw*, we have:

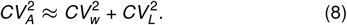

If the cell volume is simplified from (6) to 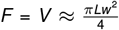, we can approximate:

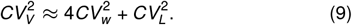

In the general case in which *A* and *V* do not scale perfectly linearly with *L*, the general formula follows (7). In SI Appendix, Section S3, through simulations, we quantify the relationship between the noises in each cell dimensions using the capsule shape finding that (9) is a reasonable approximation. Comparing the expressions (8) and (9), we conclude that the volume will be a particularly noisy variable, since the contribution of the noise in width 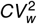 is amplified by four. Fig. 4E shows the difference in noise for cell length, cell surface area, and cell volume at different times after their most recent division. We verify the approximations in (8) and (9) concluding that:

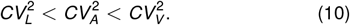

The difference between these cell dimension noises is not constant throughout the cycle. In Fig. 4, we show how the noise in the cell volume is higher for cells in early cell cycle time (newborns) and decreases as the cycle progresses. Most of the decrease in cell volume noise is related to the decrease in cell width noise, as the cell length showed a constant noise throughout the cell cycle. In SI Appendix, Section S3, we demonstrate that this decrease in volume noise is associated with the reduction of cell-width noise as cell length increases. Althought cell mass in human cells have similar noise reduction through the cell cycle [43], we ignore the origin of this regulation in our strain, but we anticipate that a further study may provide information on the mechanisms of cell surface homeostasis.

This disparity in cell size noise depending on the size proxy, especially among newborn cells (such as progenitors), is important because it defines the dynamics of population variability, as explained before. To fit our simulations, we also measured the noise of the added size 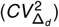. In the following section, in addition to measuring 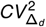, we explore how accurate the assumption of *C. glutamicum* follows the *adder* strategy.

### Size regulation of *C. glutamicum* cells

To model cell size regulation, an initial assumption is that cells grow exponentially over time. In Fig. 5A, we compare the dynamics of the cell size observed with the approximation of exponential growth. Despite the ongoing debate on the nature of *C. glutamicum* size growth [80], the results demonstrate that, for the level in detail of this study, the exponential growth approximation is a suitable choice.

**Figure 5:**
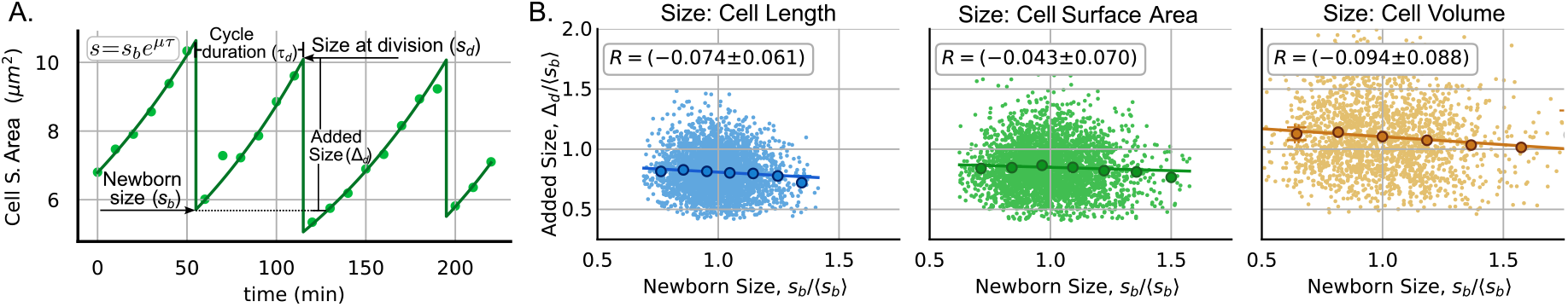
*C. glutamicum* cells control their size following the *adder* division strategy. **A**. An example of experimental trajectories of the cell surface area over time (dots). The data were fitted to exponential functions of time (continuous line). The newborn size *s*_*b*_ and the size at division *s*_*d*_ are estimated by extrapolating the fitted exponential functions of time. Other parameters such as cycle duration and added size are also represented. **B**. Division strategy represented as the relationship between the added size Δ_*d*_ and the size at the beginning of the cell cycle *s*_*b*_ for 3345 cell cycles. This strategy was presented considering cell length (blue), cell surface area (green), and cell volume (orange). The statistics of Δ_*d*_ and *s*_*b*_ are presented in Table 1. The correlation coefficient *R* between Δ_*d*_ and *s*_*b*_ is also shown with a confidence interval error of 95 % calculated using bootstrapping methods.

#### *C. glutamicum* cells divide following the *adder*

Next, we examine whether the division of *C. glutamicum* follows the *adder* division strategy. To test this hypothesis, we plot the added cell size (Δ_*d*_) against the birth size (*s*_*b*_) in Fig. 5B. The main property of the *adder* is that the added size is a variable with fixed mean independent of the size at birth [51]. This independence can be quantified by the Pearson correlation coefficient (*R*(Δ_*d*_, *s*_*b*_)) between these variables. The results confirmed that *C. glutamicum* follows an *adder* division strategy since *R* ≈ 0. To incorporate the *adder* into our simulation model, we treated the added size Δ_*d*_ as an independent random variable with the observed coefficient of variation 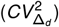. Table 2 shows the measured 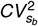 and 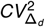 that were used in our simulations. It is worth mentioning that in a steady growing population 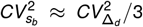 [81] which holds approximately in our observations. Finally, we also found that the added size in each cycle depends weakly on the previous added size (*R* ≈ 0, see SI Appendix, Fig. S2A. However, we observe that the added size between sister cells have a slightly positive correlation (*R* ≈ 0.17 see SI Appendix, Fig. S2B) which is similar to the observed in other bacteria [82]. We did not include these correlations to maintain the simplicity of the model.

**Table 2:** Measured parameters of cell size regulation for *C. glutamicum*. We present the mean size of the progenitor cell ⟨*s*_*b*_⟩, its variability 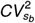, the mean added size before division ⟨Δ_*d*_⟩ and its variability 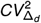. The digit in parentheses represents the amount by which the least significant digit of the value is uncertain (95% confidence interval using bootstrapping methods). For example, 1.882(6) = (1.882 ± 0.006). The analyzed dataset consists on 3345 cell cycles.

| Cell dimension | $\langle s_b \rangle$ | $CV_{s_b}^2$ | $\langle \Delta_d \rangle$ | $CV_{\Delta_d}^2$ |
| --- | --- | --- | --- | --- |
| Length ($\mu\text{m}$) | 1.882(6) | 0.0147(6) | 1.526(8) | 0.042(2) |
| S. Area ($\mu\text{m}^2$) | 5.69(2) | 0.025(1) | 4.75(3) | 0.058(2) |
| Volume ($\mu\text{m}^3$) | 1.133(7) | 0.052(2) | 1.17(1) | 0.093(4) |

### Bacterial proliferation statistics and comparison with predictions

After synchronizing the start of colony expansion from the birth of the progenitor cell, we counted the number of cells derived from each progenitor and used these data to plot Fig. 6. To compare with theory, we performed simulations using our agent-based algorithm assuming that the progenitor cell size and the added cell size during the cell cycle follow gamma distributions with the observed statistics 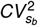 and 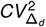 presented in Table 2. We provide more details of this algorithm in SI Appendix, Section S2.

**Figure 6:**
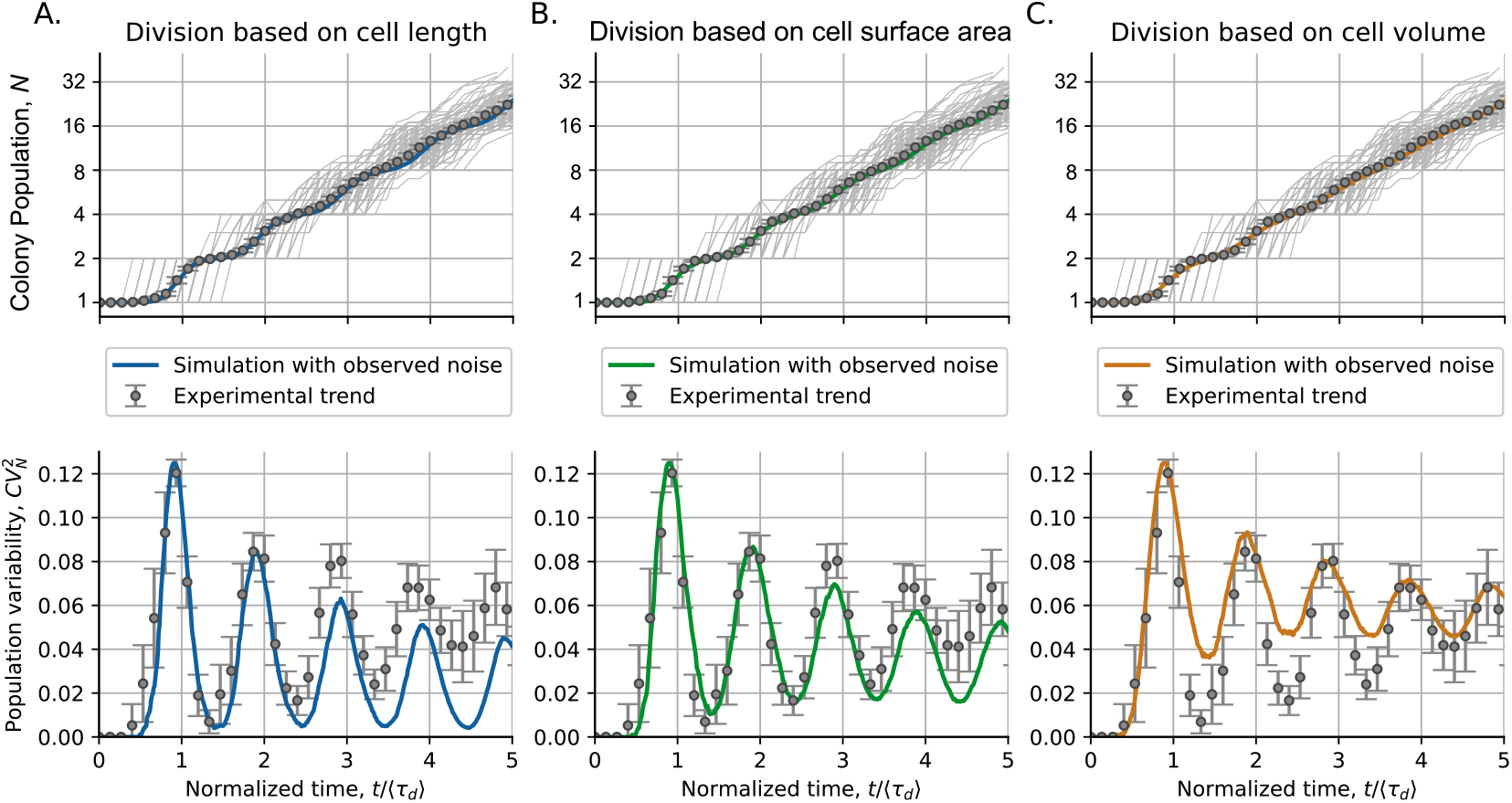
Division based on different proxies of cell size: A. length, B. surface area, C. or volume, predicts a similar mean population dynamics but different dynamics of population variability. The upper part of the Fig. shows the population in different colonies (gray lines) and their mean (error bars) compared to simulation predictions (colored lines). The bottom part shows the observed dynamics of population variability 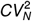 (error bars) compared to the expected dynamics from the simulations (colored lines). Simulations were performed with observed noise in the size of the progenitor cell 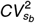 and the size of the cell added before division 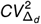. Table 2 provides the exact value of these parameters. The error bars in experiments represent the 95% confidence interval on the moments over 154 studied colonies. The mean doubling time ⟨*τ*_*d*_⟩ is 75.2 ± 0.4 min.

#### Fluctuations in colony population number aligns with the predicted if division was set by the cell surface area

Fig. 6 presents the cell proliferation statistics for the colonies of *C. glutamicum*. The statistical moments of the population distributions are compared with the results of our simulations. Since cell size can be defined using three different cell variables (length, surface area, and volume), we compare the results of the simulations using each of these types of cell size. In Fig. 6 top, we observe how the three models predict very accurate trends in the mean population ⟨*N*⟩. However, there is an appreciable difference in the prediction of trends of population variability 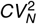 (Fig. 6 bottom).

Our model predicts that the noise in the progenitor size adjusts the basal level of oscillations (Figure 2A), while increased variability in added size reduces oscillation amplitude [20]. When analyzing the simulation based on cell length, we found that this variable shows relatively low noise (less 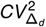 and 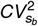) and predicts population fluctuations with stronger oscillations and a lower baseline (Fig. 6A). The model based on cell surface area predicts a trend in population variability with both the oscillation amplitude and the basal level similar to that observed experimentally (Fig. 6B). In contrast, cell volume, which has the highest variability (as explained by (10)), predicts oscillations with a smaller amplitude and a higher baseline (Fig. 6C). The close agreement between observed proliferation fluctuations and the predictions of the surface area-based model supports the conclusion that a molecular mechanism proportional to cell surface area might be the main contributor to cell division.

## Discussion

In this article, we explore how different cell division strategies predict different statistics on the colony population. Using a semi-analytical approach, we compare the *adder* strategy with the *timer* strategy. Understanding how these models describe the dynamics of population in colonies with low cell numbers is especially relevant when studying the proliferation of tumors or infections [83–85]. We expect the accuracy of these models to improve by implementing a size-dependent division.

While the dynamics of noise in colony population for *timer* division is defined by the variability of the cell cycle, in the *adder* it is defined by the variability of the progenitor cell size and the noise of the added size before division. The *timer* predicts a population variability trend that increases over time until it reaches a constant value, whereas the *adder* predicts oscillatory dynamics for the biological ranges of size control (Fig. 1C). The amplitude of these oscillations is modulated by the noise in added size: lower added size noise results in higher oscillation amplitude (Fig. 1G). In contrast, the noise in progenitor cell size determines the level around which population variability oscillates (Fig. 2A). For noiseless progenitor size, population variability increases initially, reaching a maximum during the first division, and then eventually reaches zero.

The *adder* controls cell proliferation more accurately compared to the *timer*. To have an intuitive reason behind this level of control, consider total biomass *B*, defined as the sum of all cell sizes:

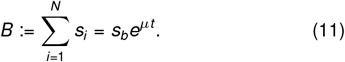

Ignoring noise in growth rate, for a given *s*_*b*_, in the SI appendix, Section S4, we show that 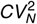 relates to cell size noise 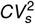 and noise in biomass 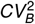 through [20]:

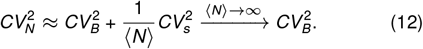

If the cell division strategy leads to a narrow cell size distribution (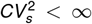 as for the *adder* [86]), 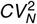 will approach 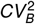. In contrast, for the *timer* division, 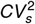 diverges [87]. This makes invalid the approximation (12). In simple words: if the cell population couples with total biomass through a cell size-based division, such as the *adder*, colony population will grow exponentially with a variability similarly to biomass.

Our model, which simplifies cell shape to a sphero-cylinder or capsule does not assume a specific proxy for cell size (volume, surface area, or length). We observe that the noise inherent in cell width causes these proxies to exhibit different levels of random variability (Fig. 4). Distinguishing the noise levels of these cell dimensions, particularly the varying noise in cell volume, surface area, and length allows us to test which dimension controls cell division by observing population growth dynamics. Although simple, our model may reveal insights into cell size regulation mechanisms for rod-shaped, exponentially growing organisms by analyzing oscillation signatures. Cells with other shapes would likely show even greater differences in the noise of their dimensions, suggesting future high-resolution imaging could enable more detailed shape analysis. Further studies with various cell strains, growth conditions, and genetic modifications can deepen our understanding of cell proliferation and division, addressing other questions in the field [50, 88].

Our findings demonstrate that the *adder* strategy provides a good approximation for cell division in *C. glutamicum* (Fig. 5). Based on this division model, we successfully predict the proliferation statistics. While the noise in progenitor size varies across the different cell size proxies we examined, our model consistently predicts similar mean population trends, though with differing population variability. Our experimental data suggest that cell division in this strain is primarily defined by cell surface area, indicating that this dimension, or a molecular factor proportional to it, is the main determinant of cell division.

The importance of the cell surface area in determining the cell cycle timing is a well-established concept. Recent discoveries in fission yeast, for example, reinforce the idea of surface area-based size control [47, 89]. These studies show that fission yeast cells halt mitotic entry until a specific surface area threshold is met. The underlying mechanisms involve proteins such as the kinase Cdr2 and the GTPase Arf6, which is crucial to anchoring the Cdr2 nodes to the cell cortex, thereby allowing surface area detection. Although our study focuses on the statistical patterns of size regulation, these identified mechanisms provide a compelling biological basis explaining why the cell surface area appears to be the growth limiting factor we observed. This context will be important for future experiments that aim to identify candidate molecular mechanisms responsible for cell size control in *C. glutamicum*.

Our model can incorporate additional factors that influence cell size regulation, such as growth rate and partitioning noises [9,81, 90]. While these variables might have minor individual effects, collectively they can significantly enhance cell proliferation variability [12]. However, incorporating all these variables is complex due to hidden correlations [91]. For example, recent studies have shown correlations between cell growth rate and size at the beginning of the cell cycle [92, 93], and between sibling cell sizes at division [94]. Other hidden variables, such as chromosome replication initiation, also play fundamental roles [13]. Understanding how all these variables correlate and contribute to cell proliferation requires more complex theoretical approaches [95, 96]. Beyond these, genetic mechanisms can impact cell proliferation by altering the growth rate or cell shape [97–101]. Other factors to consider include protein regulation effects on cell growth rate [102], different division strategies [103], and variables dependent on initial conditions [104]. However, we expect that the development of such a detailed model would add minor changes to the main conclusions of this research. This article ia an example on how the statistical properties (noise) of related variables (proliferation and cell division) can suggest underlying mechanisms providing a foundation for future research into the complex relationship between cell proliferation and size regulation in various cell types.

## Materials and Methods

### Biological preparation and medium

In this study, the bacterial strain *Corynebacterium glutamicum* ATCC 13032 was used. *C. glutamicum* was cultivated in CGXII medium [105]. CGXII medium consists of the following components per liter of demineralized water: 20 g of (NH4)2SO4, 1 g of K2HPO4, 1 g of KH2PO4, 5 g of urea, 13.25 mg of CaCl2·H2O, 0.25 g of MgSO4·7H2O, 10 mg of FeSO4·7H2O, 10 mg of MnSO4·H2O, 0.02 mg of NiCl2·6H2O, 0.313 mg of CuSO4·5H2O, 1 mg of ZnSO4·7H2O, 0.2 mg of biotin and 40 g of D-glucose. The concentration of protocatechuic acid (PCA) was 30 mg/L in the medium. For shake flask cultivation, 42 g/L MOPS buffer was added. The solutions were autoclaved or sterile filtered. Medium was adjusted to a pH of 7., All used chemicals were purchased from Carl Roth.

For preculture of *C. glutamicum*, 10 mL CGXII medium in a 100-mL baffled shake flask were inoculated from a cryostock and cultivated overnight at 30°C on a rotary shaker at 120 rpm. The preculture was transferred from a glycerol inoculated into a 100 mL shake flask with 10 mL working volume and cultivated overnight. The main culture was prepared from the preculture inoculated at an initial optical density (OD_600_) of ≈ 0.05. After reaching an OD_600_ ≈ 0.2, the culture was transferred to a 1 mL syringe and the microfluidic chip was inoculated.

### Chip design

For our microfluidic experiment, a microfluidic chip was adapted from Täuber *et al*. [74]. The chip consists on 14 arrays of cultivation chambers (80 µm x 90 µm x 650 nm) arranged in seven blocks. There are around 400 µm between each array block to separate the zones for a stable flow profile and to test five different stress duration in one experiment. The supply channels have a height of 10 µm and a width of 100 µm (See Fig. 2 in the main text).

### Chip fabrication

The microfluidic chips were fabricated using the soft lithography technique. In this process, PDMS was deposited on a silicon wafer in a ratio 10:1 between current agent and linker (Sylgard 184 Silicone Elastomer, Dow Corning Corporation, USA). After degassing the PDMS, it was baked at 80°C for 2 h. Then, the PDMS chips were cut out from the wafer and the inlets and outlets were punched (Reusable Biopsy Punch, 0.75 mm, WPI, USA). The PDMS chip and a glass slide (D 263 T eco, 39.5×34.5×0.175 mm, Schott, Germany) were cleaned three times with isopropanol and both parts were activated by O2 plasma cleaner (Femto Plasma Cleaner, Diener Electronics, Ebhausen, Germany) and assembled.

### Microfluidic single-cell cultivation

In a randomized process, the cultivation chambers were loaded with 1-5 cells. When sufficient cultivation chambers were loaded, the flow of the cell suspension was stopped and the flow of medium was started using high precision pressure pumps (Lineup series, Fluigent, Jena, Germany). A pressure of 180 and 20 mbar was set to initiate medium flow. The detailed flow profile was published in Täuber *et al*. [76].

### Live cell imaging

Live cell imaging was performed using an automated inverted microscope (Nikon Eclipse Ti2, Nikon, Germany). The microscope is placed in a cage incubator for optimal temperature control at 30 °C (Cage Incubator, OKO Touch, Okolab S.R.L., Italy). The microfluidic chip was attached to a homemade chip holder. To study the bacterial cells, a 100× oil objective (CFI P-Apo DM Lambda 100× Oil, Nikon GmbH, Germany), a DS-Qi2 camera (Nikon GmbH, Germany), and an automatic focusing system (Nikon PFS, Nikon GmbH, Germany) were used to prevent thermal drift during cultivation. 150 cultivation chambers were manually selected for each experiment using NIS Elements software (Nikon NIS Elements AR software package, Nikon GmbH, Germany). Phase-contrast images of each position were taken every 10 min with an exposure time of 50 ms and a lamp intensity of 10%.

### Simulation methods

We use Continuous-time Markov Chain Monte Carlo methods for estimating population statistics of the clonal expansion. These simulations are implemented using an agent-based algorithm in which each cell has attributes such as its own size and time to division added size since last division, time since last division and growth rate. Cells grow continuously over time following any arbitrary positively defined time derivative of the size which is followed by all agents of the simulation. The division rate of these agents is given by the division statistics studied. During division, a new agent is included in the simulation representing the additional daughter cell generated during division. The size of both cells after division will depend on the statistics of the partitioning process. We quantify the number of cells across multiple colonies and study their mean and random variability. Additional details are shown in SI Appendix, Section2 S1 and S2.

## Author contributions

- **Conceptualization:** A.G., A.S., C.N.
- **Investigation:** S.T. and L.B.
- **Data Curation:** C.N., Z.V
- **Formal Analysis:** C.N., Z.V., A.S.
- **Funding Acquisition:** A.S., A.G.
- **Methodology:** A.S, A.G.
- **Resources:** A.S, A.G.
- **Software:** C.N., Z.V.
- **Supervision:** A.S, A.G.
- **Visualization:** C.N., Z.V. S.T. and L.B.
- **Writing:** C.N., A.S.
- **Writing – Review Editing:** A.S., L.B, A.G.

## Acknowledgments

L.B. and S.T. are supported by the Joachim-Herz-Foundation (Add-on Fellowship for Interdisciplinary Life Sciences). A.S. acknowledges support from NIH-NIGMS via grant R35GM148351.

## Data Availability

Scripts for data analysis and simulations can be found at: https://zenodo.org/records/10433707.

DOI: 10.5281/zenodo.10433707.

## References

[1] César Nieto, César Vargas-García, Juan Manuel Pedraza, and Abhyudai Singh. Cell size control shapes fluctuations in colony population. In 2022 IEEE 61st Conference on Decision and Control (CDC), pages 3219–3224. IEEE, 2022.

[2] Manuel Serrano. Proliferation: the cell cycle. New Trends in Cancer for the 21st Century, pages 13–17, 2003.

[3] Marcos Malumbres and Amancio Carnero. Cell cycle deregulation: a common motif in cancer. Progress in Cell Cycle Research., 5: 5–18, 2003.

[4] Shunpei Yamauchi, Takashi Nozoe, Reiko Okura, Edo Kussell, and Yuichi Wakamoto. A unified framework for measuring selection on cellular lineages and traits. Elife, 11:e72299, 2022.

[5] Yuki Sughiyama, Tetsuya J Kobayashi, Koji Tsumura, and Kazuyuki Aihara. Pathwise thermodynamic structure in population dynamics. Physical Review E, 91(3): 032120, 2015.

[6] Yuki Sughiyama, So Nakashima, and Tetsuya J Kobayashi. Fitness response relation of a multitype age-structured population dynamics. Physical Review E, 99(1): 012413, 2019.

[7] Manuel Campos, Ivan V Surovtsev, Setsu Kato, Ahmad Paintdakhi, Bruno Beltran, Sarah E Ebmeier, and Christine Jacobs-Wagner. A constant size extension drives bacterial cell size homeostasis. Cell, 159(6): 1433–1446, 2014.

[8] John T Sauls, Dongyang Li, and Suckjoon Jun. Adder and a coarse-grained approach to cell size homeostasis in bacteria. Current Opinion in Cell Biology, 38: 38–44, 2016.

[9] César Nieto, Cesar Augusto Vargas-Garcia, and Abhyudai Singh. A moments-based analytical approach for cell size homeostasis. IEEE Control Systems Letters, 2024.

[10] Khem Raj Ghusinga, Cesar A Vargas-Garcia, and Abhyudai Singh. A mechanistic stochastic framework for regulating bacterial cell division. Scientific Reports, 6:30229, 2016.

[11] Fangwei Si, Guillaume Le Treut, John T Sauls, Stephen Vadia, Petra Anne Levin, and Suckjoon Jun. Mechanistic origin of cell-size control and homeostasis in bacteria. Current Biology, 29(11): 1760–1770, 2019.

[12] Cesar Nieto, Cesar Vargas-Garcia, and Juan M Pedraza. Continuous rate modeling of bacterial stochastic size dynamics. Physical Review E, 104(4): 044415, 2021.

[13] Prathitha Kar, Sriram Tiruvadi-Krishnan, Jaana Männik, Jaan Männik, and Ariel Amir. Using conditional independence tests to elucidate causal links in cell cycle regulation in Escherichia coli. Proceedings of the National Academy of Sciences, 120(11):e2214796120, 2023.

[14] Kaan Öcal and Michael PH Stumpf. Cell size distributions in lineages. Physical Review Research, 7(1): 013302, 2025.

[15] Arthur Genthon. Analytical cell size distribution: lineage-population bias and parameter inference. Journal of the Royal Society Interface, 19(196): 20220405, 2022.

[16] Arthur Genthon and David Lacoste. Fluctuation relations and fitness landscapes of growing cell populations. Scientific Reports, 10(1): 11889, 2020.

[17] Reinaldo García-García, Arthur Genthon, and David Lacoste. Linking lineage and population observables in biological branching processes. Physical Review E, 99(4): 042413, 2019.

[18] Ethan Levien, Jane Kondev, and Ariel Amir. The interplay of phenotypic variability and fitness in finite microbial populations. Journal of the Royal Society Interface, 17(166): 20190827, 2020.

[19] Farshid Jafarpour, Charles S Wright, Herman Gudjonson, Jedidiah Riebling, Emma Dawson, Klevin Lo, Aretha Fiebig, Sean Crosson, Aaron R Dinner, and Srividya Iyer-Biswas. Bridging the timescales of single-cell and population dynamics. Physical Review X, 8(2): 021007, 2018.

[20] César Nieto, César Augusto Vargas-García, and Abhyudai Singh. A generalized adder for cell size homeostasis: Effects on stochastic clonal proliferation. Biophysical Journal, 124(9): 1376–1386, 2025.

[21] Jie Lin and Ariel Amir. From single-cell variability to population growth. Physical Review E, 101(1): 012401, 2020.

[22] Mikihiro Hashimoto, Takashi Nozoe, Hidenori Nakaoka, Reiko Okura, Sayo Akiyoshi, Kunihiko Kaneko, Edo Kussell, and Yuichi Wakamoto. Noise-driven growth rate gain in clonal cellular populations. Proceedings of the National Academy of Sciences, 113(12): 3251–3256, 2016.

[23] Jie Lin and Ariel Amir. The effects of stochasticity at the single-cell level and cell size control on the population growth. Cell systems, 5(4): 358–367, 2017.

[24] Yaïr Hein and Farshid Jafarpour. Asymptotic decoupling of population growth rate and cell size distribution. Physical Review Research, 6(4): 043006, 2024.

[25] Daniel Taylor, Nia Verdon, Peter Lomax, Rosalind J Allen, and Simon Titmuss. Tracking the stochastic growth of bacterial populations in microfluidic droplets. Physical Biology, 19(2): 026003, 2022.

[26] Eric W Jones, Joshua Derrick, Roger M Nisbet, William B Ludington, and David A Sivak. First-passage-time statistics of growing microbial populations carry an imprint of initial conditions. Scientific Reports, 13(1): 21340, 2023.

[27] Spencer Hobson-Gutierrez and Edo Kussell. Evolutionary advantage of cell size control. Physical Review Letters, 134(11): 118401, 2025.

[28] Russell Lande. Natural selection and random genetic drift in phenotypic evolution. Evolution, pages 314–334, 1976.

[29] Sergio A Muñoz-Gómez, Gaurav Bilolikar, Jeremy G Wideman, and Kerry Geiler-Samerotte. Constructive neutral evolution 20 years later. Journal of molecular evolution, 89(3): 172–182, 2021.

[30] Thiparat Chotibut and David R Nelson. Population genetics with fluctuating population sizes. Journal of Statistical Physics, 167(3): 777–791, 2017.

[31] Médéric Diard and Wolf-Dietrich Hardt. Evolution of bacterial virulence. FEMS Microbiology Reviews, 41(5):679–697, 2017.

[32] Rishika Sen, Losiana Nayak, and Rajat Kumar De. A review on host–pathogen interactions: classification and prediction. European Journal of Clinical Microbiology & Infectious Diseases, 35(10): 1581–1599, 2016.

[33] Paras Jain, Atchuta Srinivas Duddu, and Mohit Kumar Jolly. Stochastic population dynamics of cancer stemness and adaptive response to therapies. Essays in Biochemistry, 66(4): 387–398, 2022.

[34] Bahram Houchmandzadeh. General formulation of luriadelbruck distribution of the number of mutants. Physical Review E, 92(1): 012719, 2015.

[35] Michael Saint-Antoine, Ramon Grima, and Abhyudai Singh. Fluctuation-based approaches to infer kinetics of cell-state switching. In 2022 IEEE 61st Conference on Decision and Control (CDC), pages 3878–3883. IEEE, 2022.

[36] EO Powell. Growth rate and generation time of bacteria, with special reference to continuous culture. Microbiology, 15(3): 492–511, 1956.

[37] Evgeny B Stukalin, Ivie Aifuwa, Jin Seob Kim, Denis Wirtz, and Sean X Sun. Age-dependent stochastic models for understanding population fluctuations in continuously cultured cells. Journal of the Royal Society Interface, 10(85): 20130325, 2013.

[38] Janina Bahnemann and Alexander Grünberger. Microfluidics in Biotechnology: Overview and Status Quo. Springer, 2022.

[39] Aaron M Streets and Yanyi Huang. Chip in a lab: Microfluidics for next generation life science research. Biomicrofluidics, 7(1): 011302, 2013.

[40] Ping Wang, Lydia Robert, James Pelletier, Wei Lien Dang, Francois Taddei, Andrew Wright, and Suckjoon Jun. Robust growth of Escherichia coli. Current Biology, 20(12): 1099–1103, 2010.

[41] Sattar Taheri-Araghi, Serena Bradde, John T Sauls, Norbert S Hill, Petra Anne Levin, Johan Paulsson, Massimo Vergassola, and Suckjoon Jun. Cell-size control and homeostasis in bacteria. Current Biology, 25(3): 385–391, 2015.

[42] Suckjoon Jun and Sattar Taheri-Araghi. Cell-size maintenance: universal strategy revealed. Trends in microbiology, 23(1): 4–6, 2015.

[43] Xili Liu, Jiawei Yan, and Marc W Kirschner. Cell size homeostasis is tightly controlled throughout the cell cycle. PLoS biology, 22(1):e3002453, 2024.

[44] Jaana Männik, Bryant E Walker, and Jaan Männik. Cell cycle-dependent regulation of ftsz in Escherichia coli in slow growth conditions. Molecular Microbiology, 110(6): 1030–1044, 2018.

[45] Mats Wallden, David Fange, Ebba Gregorsson Lundius, Özden Baltekin, and Johan Elf. The synchronization of replication and division cycles in individual E. coli cells. Cell, 166(3): 729–739, 2016.

[46] Miles Priestman, Philipp Thomas, Brian D Robertson, and Vahid Shahrezaei. Mycobacteria modify their cell size control under sub-optimal carbon sources. Frontiers in Cell and Developmental Biology, 5:64, 2017.

[47] Hannah E Opalko, Kristi E Miller, Hyun-Soo Kim, Cesar Augusto Vargas-Garcia, Abhyudai Singh, Michael-Christopher Keogh, and James B Moseley. Arf6 anchors cdr2 nodes at the cell cortex to control cell size at division. Journal of Cell Biology, 221(2):e202109152, 2021.

[48] Anna Knöppel, Oscar Broström, Konrad Gras, Johan Elf, and David Fange. Regulatory elements coordinating initiation of chromosome replication to the escherichia coli cell cycle. Proceedings of the National Academy of Sciences, 120(22):e2213795120, 2023.

[49] Sander K Govers, Manuel Campos, Bhavyaa Tyagi, Geraldine Laloux, and Christine Jacobs-Wagner. Apparent simplicity and emergent robustness in the control of the escherichia coli cell cycle. Cell Systems, 15(1): 19–36, 2024.

[50] César Nieto, César Augusto Vargas-García, Juan Manuel Pedraza, and Abhyudai Singh. Mechanisms of cell size regulation in slow-growing escherichia coli cells: Discriminating models beyond the adder. npj Systems Biology and Applications, 10(1): 61, 2024.

[51] César Nieto, Juan Arias-Castro, Carlos Sánchez, César Vargas-García, and Juan Manuel Pedraza. Unification of cell division control strategies through continuous rate models. Physical Review E, 101(2): 022401, 2020.

[52] Matteo Osella, Eileen Nugent, and Marco Cosentino Lagomarsino. Concerted control of Escherichia coli cell division. Proceedings of the National Academy of Sciences, 111(9): 3431–3435, 2014.

[53] Sayeh Rezaee, Cesar Nieto, Cesar Vargas-Garcia, and Abhyudai Singh. Inferring cell size control mechanisms through stochastic hybrid modeling. 2025.

[54] César Nieto, Sayeh Rezaee, Cesar Augusto Vargas-Garcia, and Abhyudai Singh. Joint distribution dynamics of cell cycle variables in exponentially-growing cells with stochastic division. In 2025 33rd Mediterranean Conference on Control and Automation (MED), pages 108–113. IEEE, 2025.

[55] Yaïr Hein and Farshid Jafarpour. Competition between transient oscillations and early stochasticity in exponentially growing populations. Physical Review Research, 6(3): 033320, 2024.

[56] César Nieto, Sergio Camilo Blanco, César Vargas-García, Abhyudai Singh, and Pedraza Juan Manuel. Pyecolib: a python library for simulating stochastic cell size dynamics. Physical Biology, 20(4): 045006, 2023.

[57] Farshid Jafarpour. Cell size regulation induces sustained oscillations in the population growth rate. Physical Review Letters, 122(11): 118101, 2019.

[58] Mattia Miotto, Simone Scalise, Marco Leonetti, Giancarlo Ruocco, Giovanna Peruzzi, and Giorgio Gosti. A size-dependent division strategy accounts for leukemia cell size heterogeneity. Communications Physics, 7(1): 248, 2024.

[59] Michal Letek, Efrén Ordóñez, José Vaquera, William Margolin, Klas Flardh, Luis M Mateos, and José A Gil. Diviva is required for polar growth in the mreb-lacking rodshaped actinomycete Corynebacterium glutamicum. Journal of Bacteriology, 190(9): 3283–3292, 2008.

[60] Erik C Hett and Eric J Rubin. Bacterial growth and cell division: a mycobacterial perspective. Microbiology and Molecular Biology Reviews, 72(1): 126–156, 2008.

[61] Handuo Shi, Yan Hu, Pascal D Odermatt, Carlos G Gonzalez, Lichao Zhang, Joshua E Elias, Fred Chang, and Kerwyn Casey Huang. Precise regulation of the relative rates of surface area and volume synthesis in bacterial cells growing in dynamic environments. Nature Communications, 12(1): 1–13, 2021.

[62] Nikola Ojkic and Shiladitya Banerjee. Bacterial cell shape control by nutrient-dependent synthesis of cell division inhibitors. Biophysical Journal, 120(11): 2079–2084, 2021.

[63] Prathitha Kar and Ariel Amir. Are cell length and volume interchangeable in cell cycle analysis? bioRxiv, pages 2024–07, 2024.

[64] Marta Ginovart, Daniel Lopez, and Joaquim Valls. Indisim, an individual-based discrete simulation model to study bacterial cultures. Journal of Theoretical Biology, 214(2): 305–319, 2002.

[65] Thomas E Gorochowski, Antoni Matyjaszkiewicz, Thomas Todd, Neeraj Oak, Kira Kowalska, Stephen Reid, Krasimira T Tsaneva-Atanasova, Nigel J Savery, Claire S Grierson, and Mario Di Bernardo. Bsim: an agent-based tool for modeling bacterial populations in systems and synthetic biology. PLoS ONE, 2012.

[66] Sivan Pearl Mizrahi, Oded Sandler, Laura Lande-Diner, Nathalie Q Balaban, and Itamar Simon. Distinguishing between stochasticity and determinism: examples from cell cycle duration variability. BioEssays, 38(1): 8–13, 2016.

[67] Niccolò Totis, César Nieto, Armin Küper, César Vargas-García, Abhyudai Singh, and Steffen Waldherr. A population-based approach to study the effects of growth and division rates on the dynamics of cell size statistics. IEEE Control Systems Letters, 5(2): 725–730, 2020.

[68] Arthur Genthon. From noisy cell size control to population growth: When variability can be beneficial. Physical Review E, 111(3): 034407, 2025.

[69] Maxime Deforet, Dave Van Ditmarsch, and Joao B Xavier. Cell-size homeostasis and the incremental rule in a bacterial pathogen. Biophysical Journal, 109(3): 521–528, 2015.

[70] Étienne Bernard, Marie Doumic, and Pierre Gabriel. Cyclic asymptotic behaviour of a population reproducing by fission into two equal parts. Kinetic & Related Models, 12(3), 2019.

[71] Judith Becker and Christoph Wittmann. Industrial microorganisms: Corynebacterium glutamicum. Industrial Biotechnology: Microorganisms, 1: 183–220, 2017.

[72] Joo-Young Lee, Yoon-Ah Na, Eungsoo Kim, Heung-Shick Lee, and Pil Kim. The actinobacterium Corynebacterium glutamicum, an industrial workhorse. Korean Society for Microbiology and Biotechnology, 2016.

[73] Jörn Kalinowski, Brigitte Bathe, Daniela Bartels, Nicole Bischoff, Michael Bott, Andreas Burkovski, Nicole Dusch, Lothar Eggeling, Bernhard J Eikmanns, Lars Gaigalat, et al. The complete Corynebacterium glutamicum atcc 13032 genome sequence and its impact on the production of l-aspartate-derived amino acids and vitamins. Journal of Biotechnology, 104(1–3): 5–25, 2003.

[74] Sarah Täuber, Corinna Golze, Phuong Ho, Eric von Lieres, and Alexander Grünberger. dmscc: a microfluidic platform for microbial single-cell cultivation of Corynebacterium glutamicum under dynamic environmental medium conditions. Lab on a Chip, 20(23): 4442–4455, 2020.

[75] Alexander Grünberger, Christopher Probst, Stefan Helfrich, Arun Nanda, Birgit Stute, Wolfgang Wiechert, Eric von Lieres, Katharina Nöh, Julia Frunzke, and Dietrich Kohlheyer. Spatiotemporal microbial single-cell analysis using a high-throughput microfluidics cultivation platform. Cytometry Part A, 87(12): 1101–1115, 2015.

[76] Sarah Täuber, Luisa Blöbaum, Volker F Wendisch, and Alexander Grünberger. Growth response and recovery of Corynebacterium glutamicum colonies on single-cell level upon defined ph stress pulses. Frontiers in Microbiology, 12:711893, 2021.

[77] Jean-Baptiste Lugagne, Haonan Lin, and Mary J Dunlop. Delta: Automated cell segmentation, tracking, and lineage reconstruction using deep learning. PLoS Computational Biology, 16(4):e1007673, 2020.

[78] Nikola Ojkic, Diana Serbanescu, and Shiladitya Banerjee. Surface-to-volume scaling and aspect ratio preservation in rod-shaped bacteria. eLife, 8:e47033, 2019.

[79] Ariel Amir. Cell size regulation in bacteria. Physical Review Letters, 112(20): 208102, 2014.

[80] Joris JB Messelink, Fabian Meyer, Marc Bramkamp, and Chase P Broedersz. Single-cell growth inference of Corynebacterium glutamicum reveals asymptotically linear growth. eLife, 10:e70106, 2021.

[81] Saurabh Modi, Cesar Augusto Vargas-Garcia, Khem Raj Ghusinga, and Abhyudai Singh. Analysis of noise mechanisms in cell-size control. Biophysical Journal, 112(11): 2408–2418, 2017.

[82] Motasem ElGamel and Andrew Mugler. Effects of molecular noise on cell size control. Physical Review Letters, 132(9): 098403, 2024.

[83] Matteo Italia, Fabio Dercole, and Roberto Lucchetti. Optimal chemotherapy counteracts cancer adaptive resistance in a cell-based, spatially-extended, evolutionary model. Physical Biology, 19(2): 026004, 2022.

[84] Andrei Kucharavy, Boris Rubinstein, Jin Zhu, and Rong Li. Robustness and evolvability of heterogeneous cell populations. Molecular Biology of the Cell, 29(11): 1400–1409, 2018.

[85] Niraj Kumar, Gwendolyn M Cramer, Seyed Alireza Zamani Dahaj, Bala Sundaram, Jonathan P Celli, and Rahul V Kulkarni. Stochastic modeling of phenotypic switching and chemoresistance in cancer cell populations. Scientific Reports, 9(1): 10845, 2019.

[86] Cesar Augusto Nieto-Acuna, Cesar Augusto Vargas-Garcia, Abhyudai Singh, and Juan Manuel Pedraza. Efficient computation of stochastic cell-size transient dynamics. BMC Bioinformatics, 20(23): 1–6, 2019.

[87] Cesar Augusto Vargas-Garcia, Mohammad Soltani, and Abhyudai Singh. Conditions for cell size homeostasis: A stochastic hybrid system approach. IEEE Life Sciences Letters, 2(4): 47–50, 2016.

[88] Guillaume Le Treut, Fangwei Si, Dongyang Li, and Suckjoon Jun. Quantitative examination of five stochastic cellcycle and cell-size control models for Escherichia coli and Bacillus subtilis. Frontiers in Microbiology, page 3278, 2021.

[89] Kristi E Miller, Cesar Vargas-Garcia, Abhyudai Singh, and James B Moseley. The fission yeast cell size control system integrates pathways measuring cell surface area, volume, and time. Current Biology, 33(16): 3312–3324, 2023.

[90] Frank J Bruggeman and Bas Teusink. Living with noise: on the propagation of noise from molecules to phenotype and fitness. Current Opinion in Systems Biology, 8: 144–150, 2018.

[91] Guillaume Witz, Erik van Nimwegen, and Thomas Julou. Initiation of chromosome replication controls both division and replication cycles in E. coli through a double-adder mechanism. eLife, 8:e48063, 2019.

[92] Maryam Kohram, Harsh Vashistha, Stanislas Leibler, BingKan Xue, and Hanna Salman. Bacterial growth control mechanisms inferred from multivariate statistical analysis of single-cell measurements. Current Biology, 31(5):955–964, 2021.

[93] Lee Susman, Maryam Kohram, Harsh Vashistha, Jeffrey T Nechleba, Hanna Salman, and Naama Brenner. Individuality and slow dynamics in bacterial growth homeostasis. Proceedings of the National Academy of Sciences, 115(25):E5679–E5687, 2018.

[94] Motasem ElGamel, Harsh Vashistha, Hanna Salman, and Andrew Mugler. Multigenerational memory in bacterial size control. Physical Review E, 108(3):L032401, 2023.

[95] Chance M Nowak, Tyler Quarton, and Leonidas Bleris. Impact of variability in cell cycle periodicity on cell population dynamics. PLoS Computational Biology, 19(6):e1011080, 2023.

[96] Liang Luo, Yang Bai, and Xiongfei Fu. Stochastic threshold in cell size control. Physical Review Research, 5(1): 013173, 2023.

[97] Jan Inge Øvrebø, Mary-Rose Bradley-Gill, Norman Zielke, Minhee Kim, Marco Marchetti, Jonathan Bohlen, Megan Lewis, Monique van Straaten, Nam-Sung Moon, and Bruce A Edgar. Translational control of e2f1 regulates the drosophila cell cycle. Proceedings of the National Academy of Sciences, 119(4):e2113704119, 2022.

[98] Evgeny Zatulovskiy, Michael C Lanz, Shuyuan Zhang, Frank McCarthy, Joshua E Elias, and Jan M Skotheim. Delineation of proteome changes driven by cell size and growth rate. Frontiers in Cell and Developmental Biology, 10:980721, 2022.

[99] Ian Jones, Lucas Dent, Tomoaki Higo, Theodoros Roumeliotis, Maria Arias Garcia, Hansa Shree, Jyoti Choudhary, Malin Pedersen, and Chris Bakal. Characterization of proteome-size scaling by integrative omics reveals mechanisms of proliferation control in cancer. Science Advances, 9(4):eadd0636, 2023.

[100] Diana Serbanescu, Nikola Ojkic, and Shiladitya Banerjee. Cellular resource allocation strategies for cell size and shape control in bacteria. The FEBS Journal, 289(24): 7891–7906, 2022.

[101] François Bertaux, Istvan T Kleijn, Samuel Marguerat, and Vahid Shahrezaei. Fission yeast obeys a linear size law under nutrient titration. microPublication Biology, 2023.

[102] Zhanhao Zhang, César Nieto, and Abhyudai Singh. Comparing negative feedback mechanisms in gene expression: From single cells to cell populations. In 2023 62nd IEEE Conference on Decision and Control (CDC), pages 3744–3749. IEEE, 2023.

[103] César Nieto, César Vargas-García, Juan Manuel Pedraza, and Abhyudai Singh. Modeling cell size control under dynamic environments. IFAC-PapersOnLine, 55(40):133–138, 2022.

[104] Chiara Enrico Bena, Marco Del Giudice, Alice Grob, Thomas Gueudré, Mattia Miotto, Dimitra Gialama, Matteo Osella, Emilia Turco, Francesca Ceroni, Andrea De Martino, et al. Initial cell density encodes proliferative potential in cancer cell populations. Scientific Reports, 11(1): 6101, 2021.

[105] Simon Unthan, Alexander Grünberger, Jan van Ooyen, Jochem Gätgens, Johanna Heinrich, Nicole Paczia, Wolfgang Wiechert, Dietrich Kohlheyer, and Stephan Noack. Beyond growth rate 0.6: What drives Corynebacterium glutamicum to higher growth rates in defined medium. Biotechnology and Bioengineering, 111(2):359–371, 2014.

