## supplementary Information for "Coupling Bacterial Cell Size Regulation with Clonal Proliferation Dynamics Reveals Cell Division Based on Surface Area"

### SI Appendix: Coupling Cell Size Regulation and Proliferation Dynamics of *C. glutamicum* Suggest Cell Division Based on Surface Area

César Nieto[a], Sarah Täuber[b,c], Luisa Blöbaum[b,c], Zahra Vahdat[a], Alexander Grünberger[b,c,d], and Abhyudai Singh[a,†]

[a]Department of Electrical and Computing Engineering, University of Delaware. Newark, DE 19716, USA
[b]CeBiTec, Bielefeld University. Bielefeld, Germany.
[c]Multiscale Bioengineering, Technical Faculty, Bielefeld University. Bielefeld, Germany.
[d]Institute of Process Engineering in Life Sciences: Microsystems in Bioprocess Engineering, Karlsruhe Institute of Technology. Karlsruhe, Germany.

#### S1 Simulation details for the *timer* division

Cell proliferation dynamics for cells dividing using the *timer* strategy were estimated using an object-oriented algorithm. The cell cycle duration is the main random variable in this process. This cell cycle duration is drawn from a gamma distribution $f_{\tau_d}$ with given mean division time $\langle\tau_d\rangle$ and cell cycle noise $CV^2_{\tau_d}$. The gamma distribution follows:

$$f_{\tau_d}(\tau; k_{\tau_d}, \theta_{\tau_d}) = \frac{1}{\Gamma(k_{\tau_d})\theta_{\tau_d}}\left(\frac{\tau}{\theta_{\tau_d}}\right)^{k_{\tau_d}-1} e^{-\frac{\tau}{\theta_{\tau_d}}}, \quad \text{(S1)}$$

with $\Gamma(.)$ being the gamma function. The parameters of $f_{\tau_d}$ can be estimated from $\langle\tau_d\rangle$ and $CV^2_{\tau_d}$ using:

$$k_{\tau_d} = \frac{1}{CV^2_{\tau_d}}, \quad \theta_{\tau_d} = \langle\tau_d\rangle CV^2_{\tau_d}. \quad \text{(S2)}$$

The main idea of the simulation is illustrated in Algorithm 1. A colony is represented by an array that initially contains one element. This element is the time to division of the first cell, which is drawn from the gamma distribution (S1). The simulation advances the time until this event occurs and then adds a new element to the array corresponding to a new cell. The time to division of both the parent and the daughter cells is reset to new random variables from the same distribution. If the array has more than one cell, the next division event is determined by the minimum time to division among all cells. The remaining times to division of the other cells are reduced by the elapsed time accordingly. Whenever there is a division, one new cell is added to the array. Cell simulation stops when the next time to division is scheduled to occur after the maximum time of the simulation. This time can be either the final time or a time for data sampling. The algorithm can resume after this stop. The implementation of the code can be found in our repository.

**Data:** $CV^2_{\tau_d}$, $\langle\tau_d\rangle$, $t_{max} > 0$
**Result:** $times = [(\tau_d)_1, (\tau_d)_2, \cdots, (\tau_d)_N]$

```
(τ_d)_1 ~ f_τd;
times = [(τ_d)_1];
τ_min = (τ_d)_1;
τ = τ_min;
if τ < t_max then
    while τ < t_max do
        m = argmin(times);
        τ_min = min(times);
        for i = 0; i < length(times); i = i + 1 do
            times[i] = times[i] − τ_min;
        end
        times[m] ~ f_τd;
        N = length(times);
        times[N] ~ f_τd;
        m = argmin(times);
        τ_min = min(times);
        τ = τ + τ_min;
    end
end
for i = 0; i < length(times); i = i + 1 do
    times[i] = times[i] − (t_max − (τ − τ_min));
end
```

**Algorithm 1:** Algorithm for the simulation of the proliferation of one colony considering the timer division strategy. Given the size mean cycle duration $\langle\tau_d\rangle$, its variability $CV^2_{\tau_d}$ and the time to end the simulation $t_{max}$, the algorithm calculates the array *times* corresponding to the time left to next division for all cells in colony. The population corresponds to the number of elements of *times*. $gamma(k_\tau, \theta_\tau)$ corresponds to a gamma distributied random variable with parameters explained in (S2).

#### S2 Simulation details for the *adder* division

The main sketch of the simulation is presented in the algorithm 2. Observe that one of the inputs of the simulation is the progenitor size $s_b$. As explained in the main text, this is a random variable. Given the variability on the progenitor cell size $CV^2_{s_b}$, we assume that the mean progenitor size satisfies $\langle s_b\rangle = \langle\Delta_d\rangle = 1$. Therefore, $s_b$ is drawn from a gamma distribution with parameters estimated from $\langle s_b\rangle$ and $CV^2_{s_b}$ from formulas similar to (S2).

In the *adder* division strategy, the added size during a cell cycle is a random independent variable. In our simulation, we consider $\Delta_d$ to be a gamma distributed variable similarly as (S1):

$$f_{\Delta_d}(y; k_{\Delta_d}, \theta_{\Delta_d}) = \frac{1}{\Gamma(k_{\Delta_d})\theta_{\Delta_d}} \left(\frac{y}{\theta_{\Delta_d}}\right)^{k-1} e^{-\frac{y}{\theta_{\Delta_d}}}, \quad \text{(S3)}$$

where the parameters can be estimated from $\langle \Delta_d \rangle$ and $CV^2_{\Delta_d}$ using:

$$k_{\Delta_d} = \frac{1}{CV^2_{\Delta_d}}, \quad \theta_{\Delta_d} = \langle \Delta_d \rangle CV^2_{\Delta_d}. \quad \text{(S4)}$$

Let $s_b$ be the size at the beginning of the cycle and $\Delta_d$ be the added size for that cycle. Thus, the size of division is given by $s_d = s_b + \Delta_d$. If we let the cell size to grow exponentially over time $s_d = s_b e^{\mu \tau_d}$ with $\mu$ being the cell growth rate, therefore, the cell cycle time $\tau_d$ is given by:

$$\tau_d = \frac{1}{\mu} \ln\left(\frac{s_b + \Delta_d}{s_b}\right). \quad \text{(S5)}$$

In addition to randomness of the size of the progenitor cell, another difference from the simulation for the *timer* division is that each cell size must be tracked. To do that, we needed an additional array that we called *sizes* in Algorithm 2. Each element is the cell size at division to the next cell splitting. Again, after each division, a new element is added to the arrays *times* and *sizes* corresponding to one cell. The implementation of the code can be found in our repository.

#### S3 Simulation of the origin of cell size scaling exponents

In the main article, we observe that the scaling exponent, the parameter that relates length to other dimensions, is greater than 1. To directly quantify the origins of this non-linearity, we analyze the dimensions of different simulated cells with capsule shape. Specifically, we investigate how much of this non-linearity arises from the capsule shape itself, how much is due to the increase in width with length, and how much results from the decrease in width noise as length increases.

##### Size scaling for cells with length-independent width

The first scenario studied consisted of estimating the scaling exponent for capsule-shaped cells with constant with. This simulation will also help us check how length scales with other cell dimensions in experiments with cell confinement. Figure S1A shows the scaling of these cell dimensions as the length increases. Each data point was generated as follows:

**a)** We consider a cell that is exponentially elongating with a random cycle timer (the time spent since the most recent division) $\tau$ that follows a uniform distribution in the interval $(0, 77)$ min. This uniform distribution of the cell cycle timer is similar to that found experimentally [1].

**b)** For each simulated cell, we generate a random cell length at birth $L_b$ from a gamma distribution with mean $\langle L_b \rangle = 1.7\mu m$ and $CV^2_{L_b} = 0.015$ with moments similar to those observed experimentally. With random generated $\tau$ and a constant growth rate $\mu = \ln(2)/77 min = 0.009\ min^{-1}$ is set such that the length doubles after $77 min$. Each cell length $L$ is calculated as follows:

$$L = L_b e^{\mu\tau}; \quad L_b \sim f_{L_b}; \quad \tau \sim U(0, 77), \quad \text{(S6)}$$

**Data:** $CV^2_{\Delta_d}$, $\langle \Delta_d \rangle$, $t_{max} > 0$, $s_b$,$\mu$
**Result:** {$times = [(\tau_d)_1, (\tau_d)_2, \cdots, (\tau_d)_N]$,
$sizes = [(s_d)_1, (s_d)_2, \cdots, (s_d)_N]$}
$\Delta_d \sim f_{\Delta_d}$;
$(\tau_d)_1 = \frac{1}{\mu} \ln\left(\frac{s_b+\Delta_d}{s_b}\right)$;
$(s_d)_1 = s_b + \Delta_d$;
$times = [(\tau_d)_1]$;
$sizes = [(s_d)_1]$;
$\tau_{min} = (\tau_d)_1$;
$\tau = \tau_{min}$;
**if** $\tau < t_{max}$ **then**
  **while** $\tau < t_{max}$ **do**
    $m = argmin(times)$;
    $\tau_{min} = min(times)$;
    **for** $i = 0;\ i < length(times);\ i = i + 1$ **do**
      $times[i] = times[i] - \tau_{min}$;
    **end**
    $s_b = sizes[m]/2$;
    $\Delta_d \sim f_{\Delta_d}$;
    $times[m] = \frac{1}{\mu} \ln\left(\frac{s_b+\Delta_d}{s_b}\right)$;
    $sizes[m] = s_b + \Delta_d$;
    $N = length(times)$;
    $\Delta_d \sim f_{\Delta_d}$;
    $times[N] = \frac{1}{\mu} \ln\left(\frac{s_b+\Delta_d}{s_b}\right)$;
    $sizes[N] = s_b + \Delta_d$;
    $m = argmin(times)$;
    $\tau_{min} = min(times)$;
    $\tau = \tau + \tau_{min}$;
  **end**
**end**
**for** $i = 0;\ i < length(times);\ i = i + 1$ **do**
  $times[i] = times[i] - (t_{max} - (\tau - \tau_{min}))$;
**end**

**Algorithm 2:** Algorithm for simulation of the proliferation of one colony using the adder division strategy. Given the mean added size $\langle \Delta_d \rangle$, its variability $CV^2_{\Delta_d}$ and the time to end the simulation $t_{max}$, the algorithm calculates the array *times* corresponding to the time left to next division for all cells in colony and the array *sizes* which is the array of sizes at division. The population is estimated as the number of elements of *times*. $gamma(k_{\Delta_d}, \theta_{\Delta_d})$ corresponds to a gamma distributied random variable with parameters explained in (S2).

where $U(0, 77)$ corresponds to the uniform distribution between 0 and 77.

**c)** Cell width $w$ was an independent random variable with normal distribution with mean $\langle w \rangle = 0.95\mu m$ and standard deviation $\sigma_w = 0.085\mu m$. This relatively low random variability justifies the use of a normal distribution.

**d)** From the random generated $w$ and $L$, the other cell dimensions were estimated using the formulas:

$$\begin{aligned} A &= \pi L w; \quad V = \pi(L-w)\left(\frac{w}{2}\right)^2 + \frac{3}{4}\pi\left(\frac{w}{2}\right)^3 \\ A_p &= w(L-w) + \pi\left(\frac{w}{2}\right)^2. \end{aligned} \tag{S7}$$

Figure S1A shows the scaling exponents of capsule-shaped cells and width with moments that are independent of cell length. Since the surface area $A$ is linear with $L$, we found an exponent of 1. However, both, the projected area $A_p$ shows an exponent of 1.09 while the volume $V$ shows an exponent of 1.33. We can also observe that the size noise between length, area, and volume is clearly different and approximately constant throughout the cell cycle. Since $CV_w^2 = (0.085)^2/(0.95)^2 = 0.008$ and $CV_L^2 = 0.015$. The noises in surface area $CV_A^2$ and volume $CV_V^2$, follow approximately equation (9) in the main text:

$$\begin{aligned} CV_A^2 &\approx CV_w^2 + CV_L^2 \approx 0.023 \\ CV_V^2 &\approx 4CV_w^2 + CV_L^2 \approx 0.047. \end{aligned} \tag{S8}$$

##### Size scaling for cell width that increases with length

Another property observed of this strain is that mean cell width increases with cell length. To estimate how the scaling exponent will change with the effects of this property, we simulate capsules with these properties. Each data point was generated following the following steps:

**a)** Cell length $L$ was generated randomly using identical procedures as the previous case (numeral a) and b)).

**b)** Given $L$, the cell width $w$ was a random variable following:

$$w = \alpha L^{\beta} + \eta; \quad \eta \sim N(0, \sigma_w), \tag{S9}$$

with $\alpha = 0.91$ and $\beta = 0.057$. $\eta$ is the noise in $w$ following a normal distribution with mean 0 and standard deviation $\sigma_w = 0.085\mu m$

**c)** From the selected $w$ and $L$, the other cell dimensions were estimated using (S7).

Figure S1B shows the cell size scaling with this non-constant width. Overall, the scaling exponents increase compared to the ones obtained with the constant-width simulation. The scale exponent between projected surface and length is 1.15, the scaling exponent between surface area and length is 0.06. Finally, the exponent between volume and simulated length is 1.45 which is very close to the ones found experimentally. The cell size noise increases slightly compared to the observed noise and is still almost constant throughout the cell cycle. However, since the variance of cell width was independent of the cell length and the mean width increases, the squared coefficient of variability of volume decreases with length and, therefore, with cell cycle time.

##### Size scaling for cell width noise that decreases with length

In the main text, we observe that the cell volume noise decreases over the cell cycle. Similarly, we note that the width variability of the cells also decreases over the cell cycle. We hypothesize that this reduction in width variability could be the cause of the reduction in volume noise along the cell cycle. To test this hypothesis, we simulate the size statistics of the cells with decreasing width variability over the cycle. We generate each data point by following these steps:

**a)** Cell length $L$ was generated randomly using identical procedures as the previous case (numeral a) and b)).

**b)** Given $L$, the cell width $w$ was a random variable following:

$$w = \alpha L^{\beta} + \eta; \quad \eta \sim N(0, \sigma_w), \tag{S10}$$

with $\alpha = 0.97$ and $\beta = 0.06$. $\eta$ is the noise in $w$ following a normal distribution with mean 0 and standard deviation depends on the length following: $\sigma_w = 0.13 - 0.015L$.

**c)** From the selected $w$ and $L$, the other cell dimensions were estimated using (S7).

Figure S1C shows the cell size scaling with this non-constant width and width noise decreasing over the cell cycle. While the noise in length and surface area show almost no variability over the cell cycle, we observe a stronger decrease on volume noise. Furthermore, the scaling exponents do not change much compared to the obtained exponents with increasing width only.

#### S4 Asymptotic behavior of the population variability

Here, we study the asymptotic limit of $CV_N^2$ and how it can be related to $CV_s^2$. Let us start with the definition of the total colony biomass:

$$B := \sum_{i=1}^{N} s_i = s_b e^{\mu t} \tag{S11}$$

We assume that the mean cell size $\langle s \rangle$ can be related to the arithmetic mean over the colony:

$$\langle s \rangle := \lim_{\langle N \rangle \to \infty} \frac{1}{N} \sum_{i=1}^{N} s_i \tag{S12}$$

##### Number and biomass asymptotic limit for the *adder* strategy

If $\langle s \rangle$ is finite, we can consider that each cell size $s_i$ can be written in terms of the mean cell size $\langle s \rangle$ and each particular fluctuation around this mean $\varepsilon_i$ as follows:

$$s_i = \langle s \rangle + \varepsilon_i. \tag{S13}$$

This means that the total biomass can be written as:

$$B := \sum_i^N s_i = N\langle s \rangle + \sum_i^N \varepsilon_i, \tag{S14}$$

Thus, the population number is expressed as:

$$N = \frac{B}{\langle s \rangle} - \frac{\sum_i^N \varepsilon_i}{\langle s \rangle}. \tag{S15}$$

Estimating the moments of (S15), we use $\langle \sum_i^N \varepsilon_i \rangle = 0$ and for the second moment:

$$\left\langle \sum_i^N \varepsilon_i \sum_j^N \varepsilon_j \right\rangle = \left\langle \sum_i^N \varepsilon_i^2 \right\rangle \approx \langle N \rangle \mathrm{var}(\varepsilon), \tag{S16}$$

in which the variance of $\varepsilon$ has the property $\mathrm{var}(\varepsilon) := \mathrm{var}(s) < \infty$, being the variance of the cell size distribution. Finally, taking the square of (S15) and taking the averages, we conclude that the moments of (S15) present the asymptotic behavior:

$$\begin{aligned} \langle N \rangle &\xrightarrow{\langle N \rangle \to \infty} \frac{B}{\langle s \rangle} \\ \langle N^2 \rangle &\xrightarrow{\langle N \rangle \to \infty} \frac{B^2}{\langle s \rangle^2} + \langle N \rangle \frac{\mathrm{var}(s)}{\langle s \rangle^2}, \end{aligned} \tag{S17}$$

which can be used for estimation of $CV_N^2$

$$CV_N^2 := \frac{\mathrm{var}(N)}{\langle N \rangle^2} \approx CV_B^2 + \frac{1}{\langle N \rangle} CV_s^2, \tag{S18}$$

in which we also used $\langle N \rangle \langle s \rangle \approx \langle B \rangle$ for a large enough population. In particular, there are two main sources of population noise, the noise in biomass that can arise from the noise in $s_b$ given by the definition (S11) or the noise in growth rate $\mu$, which was not studied in this article. The noise in the size of the cell $CV_s^2$ contributes to the transient dynamics since it is buffered by the factor $1/\langle N \rangle$.

#### References


[1] Farshid Jafarpour, Charles S Wright, Herman Gudjonson, Jedidiah Riebling, Emma Dawson, Klevin Lo, Aretha Fiebig, Sean Crosson, Aaron R Dinner, and Srividya Iyer-Biswas. Bridging the timescales of single-cell and population dynamics. *Physical Review X*, 8(2):021007, 2018.

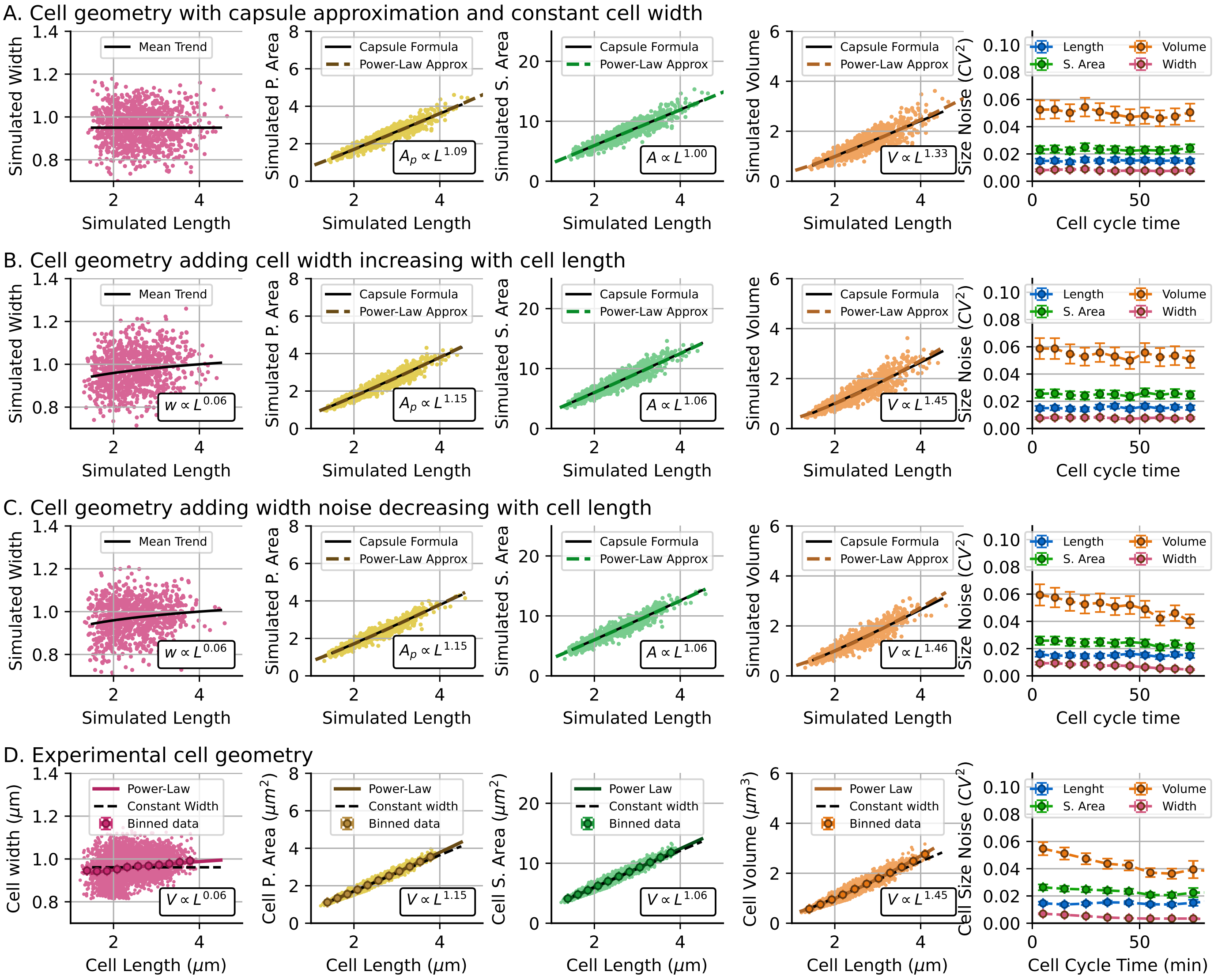


Figure S1: **Origin of the observed trends in cell size scaling** Trends of cell width (pink), cell protected area (yellow), cell surface area (green) and cell volume (orange) and their scaling exponent with cell length. Cell size noise for different cell size proxies are compares as function of the cell cycle time (time since the most recent division) **A.** Geometric properties of simulated capsules with average width independent of the length and constant noise **B.** Geometric properties of simulated capsules with a width increasing with capsule length and constant noise **C.** Geometric properties of simulated capsules with a width increasing with capsule length and width noise decreasing with length. The scatter plots show 1 000 points while the statistics was done with 10 000 data points. The simulation generates capsules with a random initial length, ($L_b$), where the mean and noise are based on experimental data. The cell cycle time, ($\tau$), is randomly drawn from a uniform distribution in the range $(0, 77)$ min. The cell length evolves according to the equation ($L = L_b e^{\mu\tau}$), where the observed growth rate is ($\mu = 0.009\ min^{-1}$). Each cell is assigned a random width, which depends on its length. Given the length and width, the remaining cell dimensions are estimated accordingly. The most likely exponent (dashed line) is determined using least-squares fitting, while the capsule model (solid line) assumes the trend of the width (function of cell length) and the length. **D.** Similar trends in cell geometry but using the experimental data (40891 points).

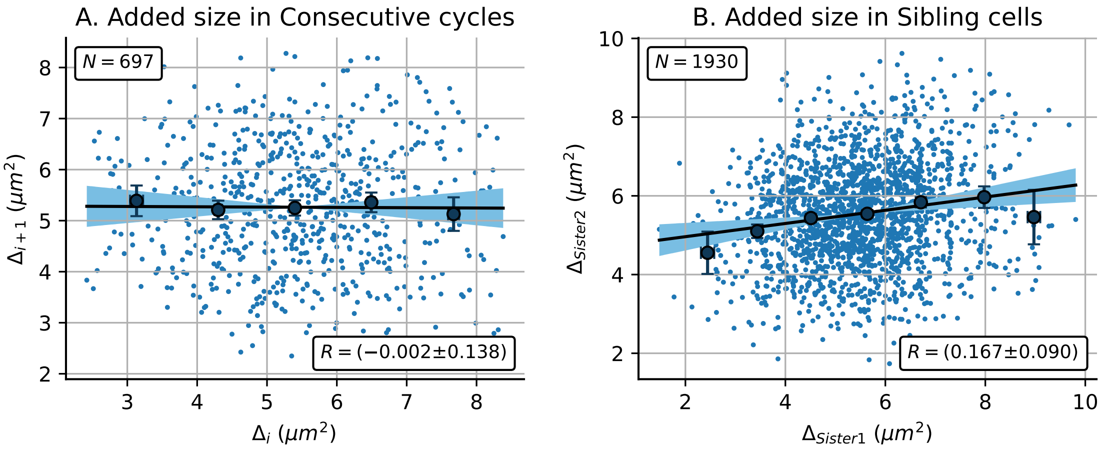


Figure S2: **Correlation of cell cycle variables between sibling cells. A.** Added size in cell cycle $i+1$ (denoted as $\Delta_{i+1}$ instead $\Delta_{d\,i+1}$ for simplicity) as a function of the added size in the previous cell cycle ($\Delta_i$). **B.** Added size of a cell sister ($\Delta_{\text{sister2}}$) as a function of the added size of the other cell sister ($\Delta_{\text{sister}_1}$) after the division that gave birth to both cells. The top left text boxes show the number of analyzed points, while the bottom right boxes show the correlation coefficients with a 95% confidence interval using bootstrapping methods.
